# Emergence of border-ownership by large-scale consistency and long-range interactions: Neuro-computational model to reflect global configurations

**DOI:** 10.1101/2021.06.17.448869

**Authors:** Naoki Kogo, Vicky Froyen

**Affiliations:** Department of Biophysics, Donders Institute of Brain, cognition and Behaviour, Radboud University, The Netherlands; Laboratory of Experimental Psychology, Brain and Cognition, University of Leuven, Belgium; Center for Cognitive Science, Rutgers University, United States

**Keywords:** figure-ground organization, border-ownership, perceptual organization, long-range neural interactions, occlusion, T-junction

## Abstract

The visual system performs remarkably well to perceive depth order of surfaces without stereo disparity, indicating the importance of figure-ground organization based on pictorial cues. To understand how figure-ground organization emerges, it is essential to investigate how the global configuration of an image is reflected. In the past, many neuro- computational models developed to reproduce figure-ground organization implemented algorithms to give a bias to convex areas. However, in certain conditions, a convex area can be perceived as a hole and a non-convex area as figural. This occurs when the surface properties of the convex area are consistent with the background and, hence, are grouped together in our perception. We argue that large-scale consistency of surface properties is reflected in the border-ownership computation. We developed a model, called DISC2, that first analyzes relationships between two border-ownership signals of all possible combinations in the image. It then enhances signals if they satisfy the following conditions: 1. the two signals fit to a convex configuration, and 2. the surface properties at the locations of the two signals are consistent. The strength of the enhancement decays with distance between the signals. The model gives extremely robust responses to various images with complexities both in shape and depth order. Furthermore, we developed an advanced version of the model (“augmented model”) where the global computation above interacts with local computation of curvilinearity, which further enhanced the robust nature of the model. The results suggest the involvement of similar computational processes in the brain for figure-ground organization.

---

Capturing the 3D structure of the surrounding environment is vital for daily life. Nevertheless, deciding the depth order of regions in 3D space involves complex computational processes. At the beginning of signal processing, information about the 3D world is underrepresented in the two dimensionally organized arrays of sensors on the retina. This underrepresentation inevitably necessitates the system to develop subjective interpretations of the environment. Despite the arbitrary nature, the human visual system performs remarkably well in judging depth order of regions even without stereo disparity cues. This indicates highly elaborate processes underlying the interpretation of depth order, or figure-ground organization, based on pictorial cues. The computational processes that give rise to figure-ground organization must have evolved in the visual system through repetitive experiences of sensory patterns and by detecting specific regularities in them. Hence, the way signals are processed must be a reflection of the way the visual system adopts statistical properties extracted from nature to aid humans to better interact with the environment. What properties exist that the visual system detects, and how are they processed? Gestalt psychologists pointed out that important cues like good continuation, symmetry, and convexity contribute to figure-ground organization. The most principal notion by the Gestalt psychologists is, however, that the computation of figure-ground organization reflects the global configuration of the image. Instead of relying on collective local properties, the computational processes capture the global structure of the image, and depth order of separate areas is computed accordingly. How is the global structure captured and reflected in the interpretation of scenes? For neuro-computational research today, this fundamental question remains to be answered.

It has been argued that the concept of border-ownership (BOWN) is key to understanding the underlying mechanisms for figure-ground organization (Koffka, 1935; Nakayama & Shimojo, 1990). At each location of a border between two segmented areas, BOWN indicates that one surface is in front of the other and that the borderline is an edge of the closer surface (the “owner” of the border). This is illustrated in Figure 1. It is assumed that there are two competing BOWN signals at each point along the borderline representing the ownerships of the two sides (Figure 1A). When an image (as shown in Figure 1B) is presented, figure-ground organization is established so that the black disk is perceived to be on top of the gray surface that continues behind it (Figure 1C). The border is an edge of the black surface, and hence is “owned” by the black side. In other words, the borderline is “intrinsic” for the black surface but “extrinsic” for the gray surface (Nakayama et al., 1989). Importantly, BOWN signals reflect global configurations of the image as illustrated in Figures 1D and 1E. In Figure 1D, the gray rectangle is perceived to be on top of the black surface which corresponds to the ownership of the central border by the gray rectangle. By removing a small black rectangular area at the bottom (solid black arrow in Figure 1D) and adding a gray rectangle of the same size on top (solid black arrow in Figure 1E), the black rectangle is now perceived to occlude the gray rectangle (Zhou et al., 2000). This coincides with the reversal of the BOWN signal at the central border. The reversal is not due to the local properties around the border as they are exactly the same in the two images, but rather due to the change of the remote components (the small rectangles). This indicates that, while the BOWN signal is assigned locally, the signal itself reflects global configurations.

**Figure 1:**
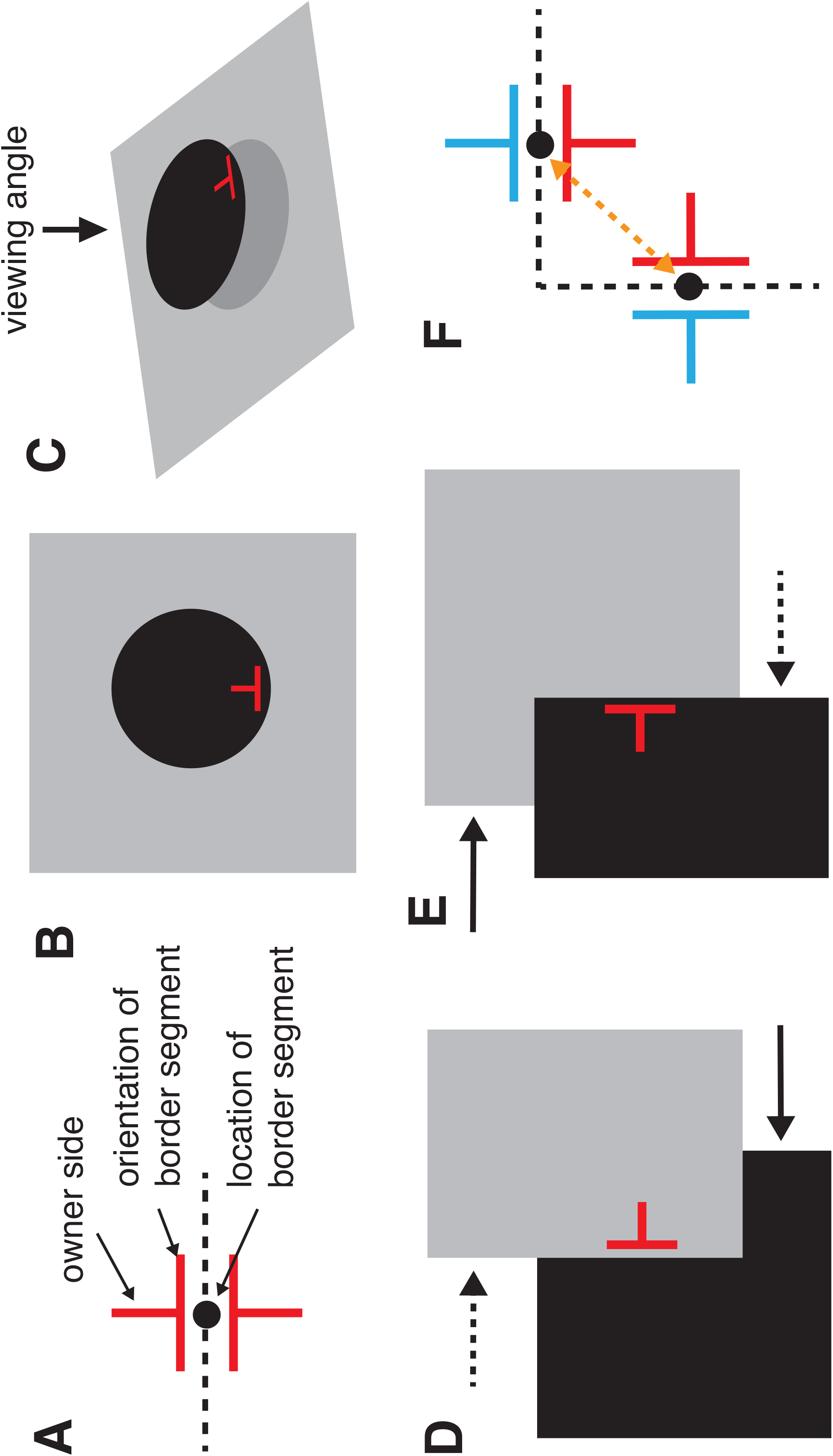
(A) Competing BOWN signals at a point on a borderline (black dot). The T-shaped symbol is used as a symbol of BOWN signal in this paper. The fin perpendicular to the border segment depicts the owner side and its length indicates the strength of the ownership. (B) An image perceived as a black circle on gray background, with an example BOWN signal in red. (C) The 3D interpretation of the image shown in (B). (D, E) Illustration of the influence of global configuration on a local BOWN signal (red). Modified from Zhou et al., 2000 (see the main text for details). (F) Illustration of four competing BOWN signals present at an L- junction.

Importantly, neurons that show tuning for BOWN (BO cells) were found in the lower level visual cortex of monkeys (Zhou, Friedman, & von der Heydt, 2000). Follow-up studies have shown that these neurons are tuned to stereo disparities as well (Qiu & von der Heydt, 2005), that the BOWN sensitive response has a short onset latency and hence may involve a feedback circuit (Craft et al., 2007; Sugihara et al., 2011; Zhou et al., 2000), that the neural circuit for the BOWN computation may be involved in selective attention (Mihalas, Dong, von der Heydt, & Niebur, 2011; Qiu, Sugihara, & von der Heydt, 2007), and that BO cells show BOWN sensitivities to objects in natural images and illusory contours (Hesse & Tsao, 2016). An fMRI study has also shown an evidence for the computation of BOWN in human visual cortex (Fang et al., 2009). These data indicate that BOWN computation processes are likely playing important roles mediating signal processes between lower-level feature detections and higher-level global properties.

For the BOWN signals to reflect global configurations of given images, BOWN computation mechanisms must be made such that BO cells are coherently activated based on the global structure of the image. How is this done? It is key to note that large-scale *relationships between the local elements* must be playing an important role in computational processes. It is possible that when the signals are coherent, the activities of BO cells are enhanced and, through long-range interactions, the specific patterns of activities of BO cells emerge. In fact, it has been shown that the enhancement of the activities of BO cells is observed when a single edge element (instead of a complete enclosed edge) is given at a distance, if the edge element is consistent with the edge element given at the receptive field of the recorded BO cell (Zhang & von der Heydt, 2010). The importance of this observation cannot be emphasized enough: It indicates that for the interactions between BO cells to occur, it is not necessary that the border elements in the input image are connected. Thus, long-range interaction mechanisms have to be assumed. The next question is, then, to identify which neural mechanisms mediate long-range interactions.

The data by Zhang et al. is not consistent with the idea that neighboring BOWN signals interact and that final outcomes emerge as the result of the horizontal propagations of signals along the borders. Furthermore, the very short onset latency of the BOWN-sensitive component of BO cells’ response cannot be explained by horizontal connections with their slow conduction velocity (Craft et al., 2007; Sugihara et al., 2011; Zhou et al., 2000). These physiological data give the key constraints to model the underlying mechanisms of BOWN computation and, consequently, it has been suggested that the computation is done by inter-areal feedforward/feedback connections mediating long-range interactions between BOWN signals (Craft et al., 2007; Zhou et al., 2000). If the long-range interactions as such underlie the coherent activations of BO cells, the interactions must be done according to specific rules based on the specific relationships between the two signals. Hence, to understand the BOWN computation mechanisms it is essential to investigate based on what rules the interactions are made. The results of these specific interactions should also in turn correspond to the specific figure-ground organization reflecting the global structure of the image. One such “rule of interaction” is what we call a “convexity bias”, which has been implemented in many neuro- computational models.

There are large number of computational models that simulated figure-ground organization (Baek & Sajda, 2005; Craft et al., 2007; Finkel & Sajda, 1992; Froyen et al., 2010; Goldreich & Peterson, 2012; Grossberg & Yazdanbakhsh, 2005; Jehee et al., 2007; Kienker et al., 1986; Kikuchi & Akashi, 2001; Kikuchi & Fukushima, 2003; Kogo et al., 2010; Layton et al., 2012; Roelfsema et al., 2002; Sajda & Finkel, 1995; Sakai et al., 2012; Sakai & Nishimura, 2006; Vecera & O’Reilly, 1998; Zhaoping, 2005). Despite the differences of the algorithms and configural cues they implemented, one commonality may be found among them. First, these algorithms assume competitions between abutting segmented areas for figure assignment, or competitions between BOWN signals with opposite directions of ownerships. Second, the algorithms give bias to the competition in such a way that the side of a convex area becomes the winner. In other words, the algorithms are made so that the inside of an enclosed area becomes the figure. For example, the algorithm implemented in the computation of BOWN or edge assignment in many models (Craft et al., 2007; Finkel & Sajda, 1992; Hu et al., 2019; Jehee et al., 2007; Kienker et al., 1986; Kikuchi & Akashi, 2001; Kikuchi & Fukushima, 2003; Kogo et al., 2010; Layton et al., 2012; Sajda & Finkel, 1995; Vecera & O’Reilly, 1998; Zhaoping, 2005) can be generalized as follows: Assume that there is an L-junction as shown in Figure 1F where four possible BOWN signals are indicated. The algorithm is made so that the BOWN signals interact either directly, or via a hierarchical feedback circuit. In these interactions, the two signals that indicate ownership of the inside of the L-junction (“inward ownership”, red) are enhanced. This enhancement is made to be stronger than the interaction

(if any) of the other two BOWN signals with “outward ownerships” (blue). This algorithm can be applied to junctions of any angles between the two border segments, and bias is given to the BOWN signal pair with the smaller angle side. If there are collections of junctions with infinitesimally small border segments and one is connected next to the other, it forms a curved border. Assume a set of border segments from an object where they collectively form an enclosed border. In a simple shape such as a circle, a pair of two inward BOWN signals at two neighboring points always forms a smaller angle than its outward counterpart. Even if the enclosed borderline forms a complex shape, the sum of the angles of inward BOWN signal pairs is smaller than outward BOWN signal pairs due to the geometrical constraint of enclosure (Feldman & Singh, 2005). Hence, with the algorithm that gives bias to the side with smaller angles, the balance of the competition is shifted towards the inside of the enclosed area. Consequently, the interior of the enclosure becomes the owner of the border, indicating that the enclosed convex area is a figure. Hereafter, we call this approach the “convexity bias” in the computational process of ownership: at each location along a border, a bias is given to the two competing BOWN signals so that the one on the side of the convex area becomes stronger.^1^ This convexity bias algorithm results in a model response similar to the convexity preference observed in human vision. This is the essential reason why these models worked well (Jacobs, 1996; Liu, Jacobs, & Basri, 1999; Rubin, 1921; Kanizsa & Gerbino, 1976).

However, it is not always the case that convex areas are perceived as figures. In Figure 2A, the large gray disk is perceived as being on top of the textured background and the small black disk is perceived to be on top of the large disk. In contrast, the central area in Figure 2B tends to be perceived as a hole, due to the texture being consistent with the background (Bertamini, 2006; Bertamini & Hulleman, 2006; Yin et al., 1996, 2000). This suggests that the consistency of surface properties between separated surfaces play a fundamental role in figure-ground perception. Hence, any computational models that are solely based on border maps of images are bound to fail to reproduce human perception because the border maps of the two images in Figure 2 are exactly same (Figure 2C). This is a key observation that has a significant implication in modeling figure-ground computation. It is clear that on top of the geometrical layout of border signals, a model has to reflect the consistency of surface properties. This is the motivation of the development of the model reported here.

**Figure 2:**
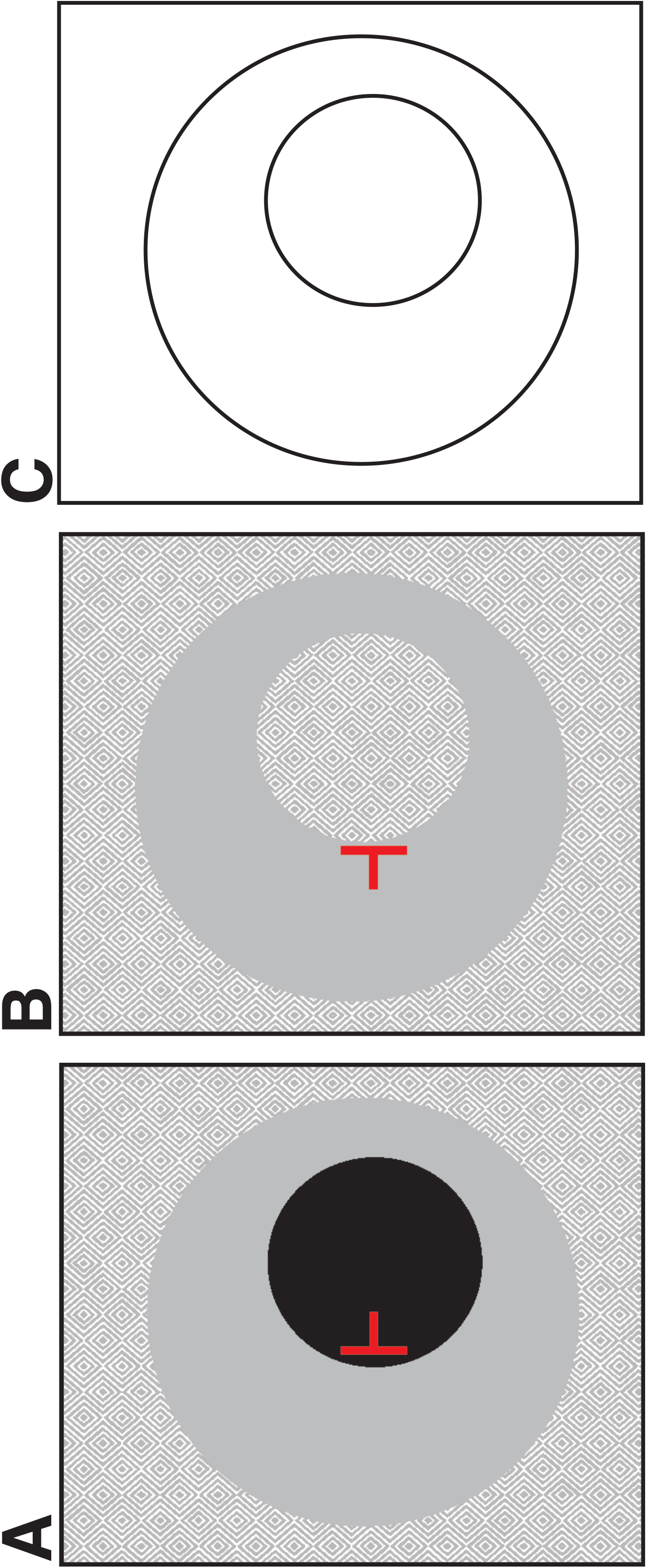
Illustration of dependency of perception of convex areas on surface consistency. (A) The central dark brown area is perceived as on top of the light gray area and hence BOWN of the central circular border is inward (red). (B) The central area is perceived as a hole and BOWN of the central circular border is outward (red). (C and D) The border maps of the two images are identical.

The necessity of reflecting the large-scale consistency of surface properties in the computation of border-ownership has been largely ignored except for a model by Zhaoping (Zhaoping, 2005). In her paper to report the biologically plausible model of border-ownership neurons, she reported a second version of the model called an “augmented model”. In this model, she reflected the data by Zhou et al. (2000) that reported that a large number of BO cells are also sensitive to a particular contrast polarity. When a luminance border is presented to the neuron’s receptive field (and if the contrast polarity matches its preferred polarity), the neuron showed BOWN tuning; however, if the polarity was reversed, the tuning disappeared (Figures 6 and 7 of Zhou et al., 2000). Zhaoping’s insight in developing the augmented model was that this sensitivity to contrast polarity aids in the detection of the consistency of surface properties and that the neural signals from BO cells are grouped (and enhanced) accordingly.

One of the authors of this paper developed a model for BOWN computation to explain illusory surface perception (DISC model, Kogo et al., 2010). Instead of implementing physiological model neurons representing BOWN (as in Zhaoping’s model), BOWN was directly computed based on the relationships between BOWN signals. While the model successfully reproduced the illusory surface perception responding to the global configurations of the image, the algorithm did not reflect the surface properties as in the augmented version of Zhaoping’s model. Therefore, the previous DISC model would not be capable of reproducing the differential perceptions of the examples shown in Figure 2. We have now developed a new version of the DISC model (called “DISC2”) with the implementation of an algorithm that reflects the consistency of surface properties. To investigate the potential of the approach by Zhaoping, we expanded the original model. First, we implemented an algorithm to detect orientations of boundaries at each location so that any arbitrary shapes can be tested by the model. This feature is essential to investigate responses of the model to key images that have important implications to perceptual organization (models by Zhaoping (2005) and Kogo et al. (2010) both limited detection of orientations of boundaries only to vertical and horizontal orientations). Second, we implemented an algorithm to make BOWN signals interact only when the surface properties at the border segments are consistent. By doing so, we demonstrate that our model shows extremely robust responses to images with complex depth configurations. In addition, we developed a two-layered model called the “augmented DISC2”, where local and global properties are computed interactively. In the augmented model, the global computation of DISC2 is combined with a local curvilinearity-based BOWN computation. This further improved the robustness of the model. These models show unprecedented robust responses, suggesting the biological plausibility of the computational principles implemented to them.

## Model

The input image is pre-processed in order to compute border nodes, and assign two competing BOWN signals at each node (for pre-processing details see Supplemental Material S1). At the initial condition of the model (before the first iteration of the algorithm), equal amounts of BOWN values (0.5) are given to all BOWN signals. Next, the relationships of all possible combinations of BOWN signals are considered and “consistent” BOWN signals (see below) interact with each other in order to compute BOWN values. The BOWN value of an individual BOWN signal reflects how many other BOWN signals are consistent with the BOWN signal in question. Consistency of two BOWN signals is determined by the two consistency constraints described below. Furthermore, a distance factor determines the strength of the interaction between consistent BOWN signals depending on the distance between them. The results of the computations are reflected in a consistency weight (more details below). The average of all BOWN values weighted by the consistency weights is added to the BOWN value from the previous iteration, which in turn constitutes the new BOWN values. These steps are then repeated until convergence. There are no specific L- or T- junction detection algorithms implemented. Algorithms for texture-based segmentation and for disentanglement of illumination and reflectance from luminance are not implemented in pre-processing. Hence, the BOWN computation is for segmented areas based on the (achromatic) colors of surfaces. The model is not made to construct 3D curvature of surfaces (volume) as it is likely to be computed by separate mechanisms. This is not within the scope of the paper (see the discussion).

We give a more detailed overview of the approach below.

## Conventions

In this paper, BOWN signals are expressed by a T-shaped symbol with a line segment (“*fin*”) that is a normal of a border segment (perpendicular to the local orientation of the border) (Figure 1). The length of the *fin* expresses the strength of the ownership. The direction of a BOWN signal is expressed by the direction of the fin in the algorithm. Angle 0^°^ of the direction corresponds to the “East” direction, and direction angles are measured counter-clockwise in a range of [-180^°^, 180^°^].

## Consistency Constraints and Detection

The computation of the consistency weight consists of three processes. The combination of the results of these three processes determines the relationship between two BOWN signals, and determines each BOWN signal’s “contribution” to another BOWN signal. First, the geometrical relationship between the two BOWN signals has to satisfy the condition of convexity. Second, the surface properties of two abutting areas that constitute the border have to be consistent at the locations of the two BOWN signals. Lastly, the strength of the interaction decays exponentially with increasing distance between the two.

### Convexity Constraint and Detection

If two BOWN signals satisfy the convexity condition, they are considered to be geometrically “consistent” with each other. To determine this, the angles that indicate geometric relationships between the two BOWN signals are computed. The first angle, *θ*, indicates the angle measured from the direction of the first BOWN signal to the line drawn between the two. The second angle, *φ*, indicates the angle measured from the direction of the second BOWN signal to the line drawn between the two (Figure 3). The relationship can be categorized by dividing the angle values into distinct classes: less than -90^°^, -90^°^, between -90^°^ and 90^°^, 90^°^, and larger than 90^°^. When the two BOWN signals have the same directions and are aligned with the second BOWN signal being on the either sides of the first BOWN signal (*θ* and *φ* values are either -90^°^ and 90^°^ or 90^°^ and -90^°^), they are *collinear* (see Figure 3A). This collinear condition is considered to be a special case of the convexity condition and the two BOWN signals are considered to be geometrically consistent. Note that collinearity means not only are they aligned, but also their directions of BOWN indicate the same side of ownership (when the directions are opposite, they are not considered to satisfy the condition). When the two angles are both between -90^°^ and 90^°^, they are considered to be in a *convex relationship* and the two BOWN signals are considered to be geometrically consistent. An example of this relationship is shown in Figure 3B: the fins in the pair point toward each other and, in this condition, one can imagine a convex shape fitting to the two BOWN signals. On the contrary, if the two angles are either lower than 90^°^ or higher than -90^°^, it creates a *concave* relationship (see Figure 3C), and they are considered to be inconsistent (Kogo & van Ee, 2015). In all other cases, the pair is considered to be *inconsistent*. Figure 3D illustrates a field of BOWN signals in relation to a given BOWN signal in the center (green). Centers of individual wheels indicate the location of BOWN signals. The red and blue rim indicate the ranges of directions of consistent (red) and inconsistent (blue) BOWN signals (with the spokes indicating example directions of BOWN signals). Some examples of BOWN signals consistent with the center BOWN signal are shown (purple). In what follows, convexity consistency of each pair of BOWN signals, with identifier indices *i* and *j*, will be indicated as *CC*(*i*, *j*), where,

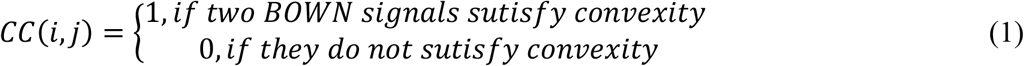

**Figure 3:**
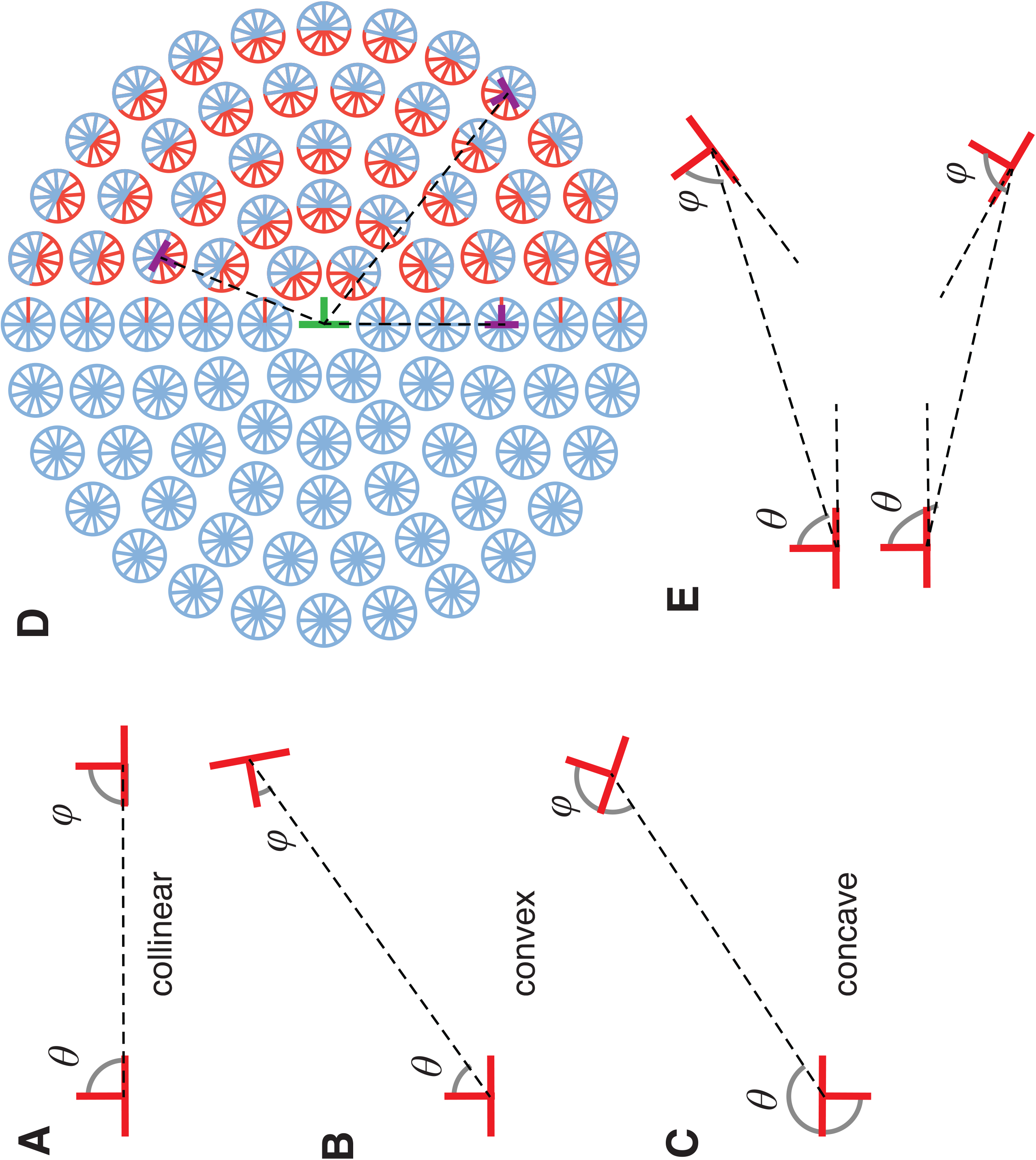
Illustration of the relationships of two BOWN signals that satisfy “convexity” condition. (A to C): The angles *θ* and *φ* indicate the angle measured from the direction of the first and second BOWN signals, respectively, to the line drawn between the two. Given these angles, several geometrical relationships emerge: collinear (A), convex (B), concave (C) and other cases (not shown). Of these, “collinear” and “convex” are consistent with convexity constraint and all other relationships are not (see the main text). (D): A field of BOWN signals in relation to a BOWN signal in the center (green). Centers of individual wheels indicate the location of BOWN signals. The red and blue rim indicate the ranges of directions of consistent (red) and inconsistent (blue) BOWN signals (with the spokes indicating example directions of BOWN signals). Examples of BOWN signals consistent with the center BOWN signal are shown (purple). (E): If border segments of two BOWN signals are curvilinearly aligned and the ownerships are on the same side, the BOWN signals are considered as curvilinear. This constraint is used for the augmented DISC2 model.

### Surface Consistency Constraint and Detection

As explained in the introduction, consistency of a surface property needs to be detected. The following observation indeed indicates that implementation of a surface consistency constraint should help BOWN computation. The central area on the image in Figure 4A has a different gray scale from the rest. In contrast, the central area in the image in Figure 4B has the same gray scale value as the background. Due to the inconsistency and consistency of the surface property with the background, respectively, the central area is perceived to be a figure on top of the larger surface in Figure 4A, and as a hole in Figure 4B (as discussed for the images in Figure 2). BOWN signals in several locations are shown in Figures 4A and B (with *i* and *o* in subscript indicating inward and outward directions of BOWNs from the center of the image, respectively). Consider the pair of BOWN signals at the bottom, *B*_2*o*_ and *B*_3*i*_ (blue). In Figure 4A, although the two BOWN signals follow the convexity condition described above, the colors at the boundaries are different: *B*_2*o*_ has 0.0 and 0.5 on the owned side (the side where the BOWN signal indicates the ownership) and the un-owned side (Figure 4A2), respectively, while *B*_3*i*_ has 0.0 and 0.8 (Figure 4A3). Because of this inconsistency in the surface property, they are considered to be an “inconsistent pair” in our model. On the other hand, the BOWN signals with inward directions for the border of the central area (*B*_1*i*_ and *B*_2*i*_, red) are consistent in the colors on both owned and un-owned sides (Figure 4A1 and 4A2). Hence, the inward BOWN signals would enhance each other and the output of the model would indicate the inside of the central area as the figural side. However, the condition is different in Figure 4B. *B*_2*o*_ and *B*_3*i*_ has the same colors on both owned and un- owned sides, 0.0 and 0.8, respectively. There are large numbers of inward BOWN signals at the outer border (e.g. *B*_3*i*_) consistent with the outward BOWN signals of the inner border (e.g. *B*_2*o*_). Hence, when the consistent pairs enhance each other, the balance of the competition for the ownership between the two BOWN signals at the inner border (*B*_2*i*_ and *B*_2*o*_) is shifted to the outward direction significantly.

**Figure 4:**
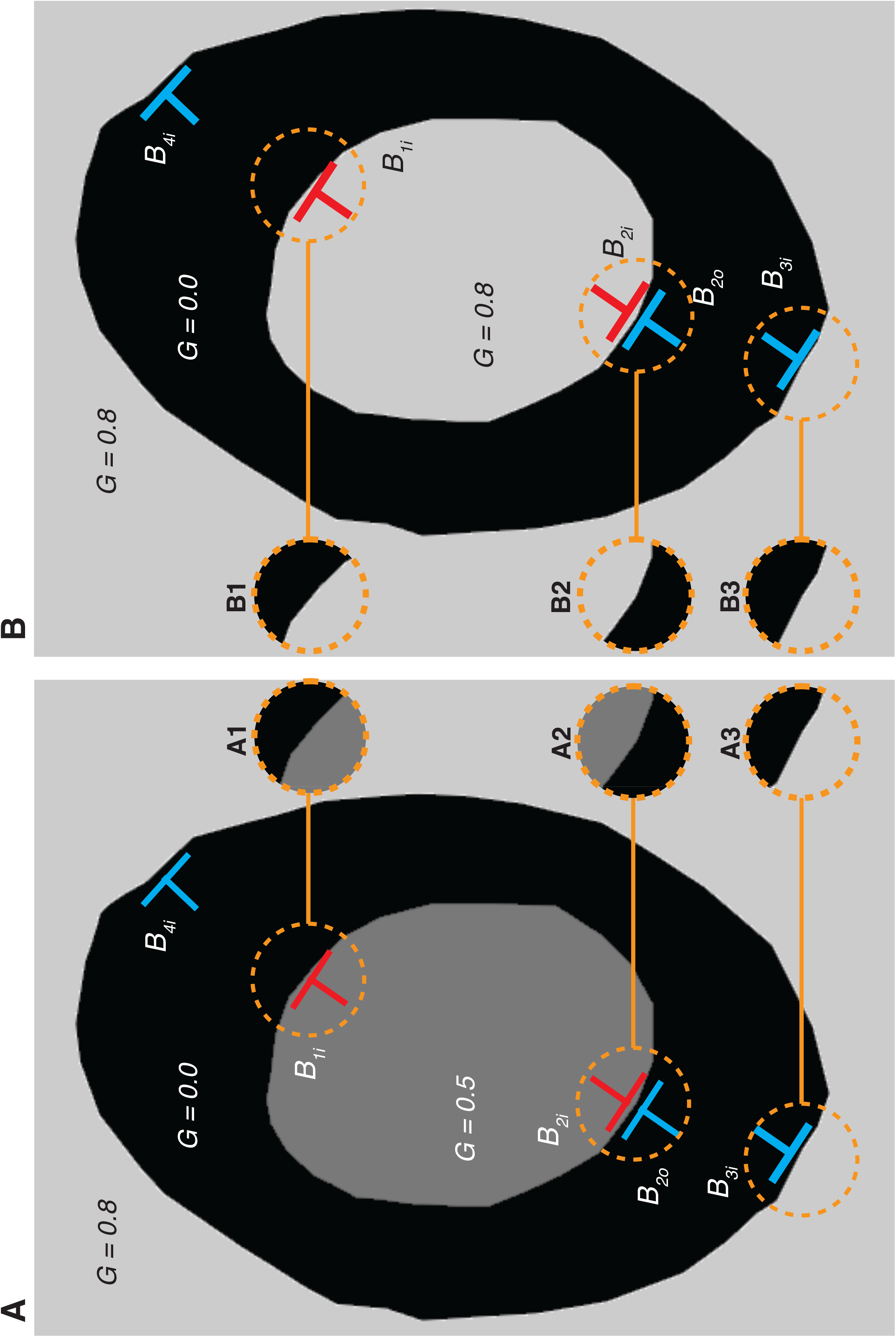
Illustration of surface consistency detection approach. The orange circles capture the gray-scale values on the owned and un-owned side of each BOWN signal (A1-A3, B1-B3). *G* indicates the gray-scale values of each area in scale of 0 to 1.0. In the model, a pair of BOWN signals is considered to have surface consistency if the gray scale values on both the owned and un-owned side correspond between the two BOWN signals. For example, *B*_2*o*_ and *B*_3*i*_ in (A) are not consistent (see A2 and A3), while the same signals in (B) are consistent, (see B2 and B3).

The surface consistency constraint here is formalized as follows. At each location of the BOWN signal, the gray scale of the surface on the owned side and the un-owned side are detected and defined as *C*_*own*_ and *C*_*unown*_. Surface consistency is then defined as follows:

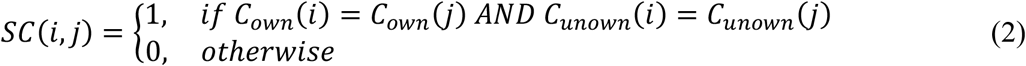

### Distance Decay Factor

Lastly, the strength of the interaction between the two BOWN signals decays exponentially with the distance between them:

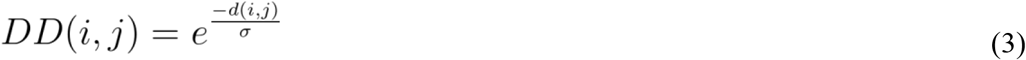

where *d*(*i*, *j*) is the distance between BOWN signals *i* and *j*. σ is a free parameter, the space constant, of the model. In this way, BOWN signals that are close to the target BOWN signal exert more influence than BOWN signals that are farther away.

In the model, only the pairs that satisfy both convexity and surface consistency conditions make contributions to the BOWN computation. This AND operation is implemented by simply multiplying both consistency conditions. Finally, we multiply this result with the distance decay function, to yield the final consistency weight.

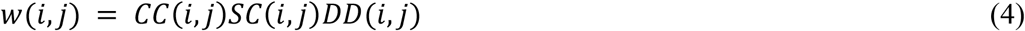

## Interaction

BOWN values are computed iteratively. Here we use the symbol *B*^*t*^(*i*) as the BOWN signal with identifier index *i* at iteration *t*. Note that at each location, there are two competing BOWN signals. Hence, for *N* border nodes, there are 2*N* BOWN signals, and the index *i* ranges from 1 to 2*N*. In the initial condition, all BOWN signals have the value of 0.5. The computation is done by averaging all other BOWN values, *B*(*n* ≠ *i*), weighted by their corresponding consistency weights *w(i,n)*, and adding it to each *B*^*t*−1^(*i*) from the previous iteration (*t* − 1) as follows^2^:

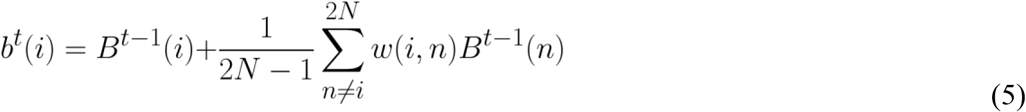

The resulting values, *b*^*t*^(*i*), are then normalized between competing signals at each location as follows:

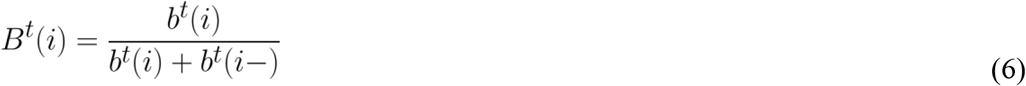

Here *i-* stands for the identity of the opposing BOWN signal at the same location as *b(i)*. This normalization is, in effect, an operationalization of mutual inhibition between two competing BOWN signals (Zhaoping, 2005). The values of *B*(*i*) (ranged from 0 to 1.0) can be interpreted as the probability that the side that the signal represents is the owner of the border at the location. The model iterates until the criterion of equilibrium is met:

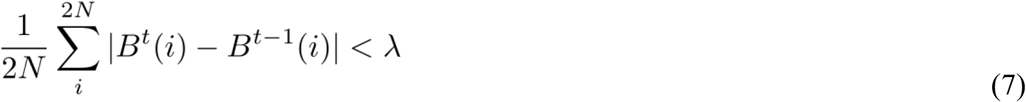

In other words, the average of the difference of all *B*(*i*) at the previous iteration and the current iteration has to be smaller than a criterion value, *λ*.

## Augmented model

In addition, we further expanded the model to solve a problem (described in section “An example with an inconsistent response” in Results). To distinguish the two models, the expanded model is called “augmented DISC2” while the model described above is called “standard DISC2”. The augmented model has two layers to compute local and global properties. The global computation is exactly the same as the standard model described above. The local computation detects curvilinearity between neighboring border segments and the consistency of surface property only on the owner-side. The two computations are combined to determine the BOWN signals. The rationale of implementing this additional step is discussed in detail in the section “Augmented model” in Results.

The “curvilinearity of BOWN signals” is defined as follows: the border segments of the two BOWN signals are curvilinearly aligned, their orientation difference is less than a maximal value, *ɛ*, and the two BOWN signals indicate the same side of ownership. Here, ownership is considered as “same side” if a line drawn between the two BOWN signals belongs to either owned side or un-owned side of both of the BOWN signals (Figure 3E top and bottom, respectively). Consistency based on the curvilinearity, is defined as follows.

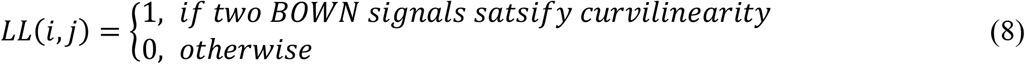

In the model, *ɛ*is set to 40°.

The surface consistency constraint for the local BOWN computation, *SC_LCL_*, is that gray scale of the surface only on the owned side *C*_*own*_ has to be consistent as follows:

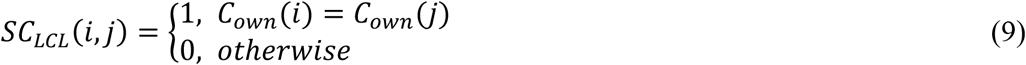

The distance decay factor, *DD_LCL_*, is the same as the one for the global computation, except that the distance between the two BOWN signals has to be shorter than *d_max_*.

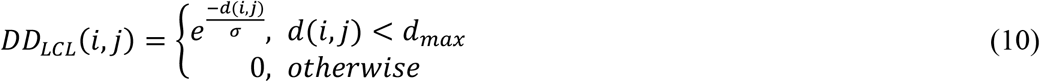

In the model, *d_max_* is set at 100.

The consistency weight of the local computation is the product of these three factors.

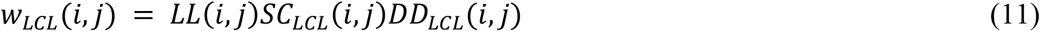

The BOWN signals by local computation, and the ones by global computation, are added at each iteration as follows.

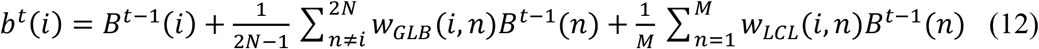

Here, the second term is for the global computation with *w_GLB_* defined by Eq. 3 and the third term is for the local computation. The normalization of *b^t^* is applied at each location as described in Eq. 6. The improvement of the model’s performance by this algorithm is described in details in the section “Augmented model” in Results.

The MATLAB code of the model is available for further testing (https://github.com/vickyf/disc2).

## Results

Here we show results from the standard model, DISC2, applied to a variety of images, ranging from simple isolated shapes to complex images that evoke perception of occlusions and holes. The size of all images used below was 800 pixels by 800 pixels. We set the space constant to *σ* = 200 which gave a smooth decrease of distance factor (*DD*) within a distance range of [0, 800]. The convergence criterion value, λ, is set to 1e^-5^. Later, we show the results from the augmented version of the model and discuss improvement of the responses.

### Examples: basic responses

First, we describe the response of the model to a simple image, an oval (Figure 5A). Figure 5C shows the initial condition of the computation, after pre-processing the image, with the detected border nodes (black dots along the border) and the initial BOWN signals (black lines) at each node. The length of each BOWN signal indicates the strength of the BOWN signal. Note that there are two BOWN signals at each node, indicating opposite owner sides. Also note that for visualization purposes, only a randomly sampled set of BOWN signals is plotted due to the high density of border nodes. Before iteration starts, the two competing BOWN signals at individual border nodes have the same strength (0.5). When iteration starts, the BOWN signals in the inward direction become quickly dominant due to the convexity of the shape ([suppl movie 1](https://www.dropbox.com/s/7c5unvyleq5dzoq/movie_suppl_1_ovalNew_half_opt.gif?dl=0)). Figure 5D and 5E shows the BOWN signals at iteration 5 and at equilibrium (iteration=46), respectively. The red and blue lines indicate BOWN signals with more and less dominance, respectively (i.e. red when BOWN signal > 0.5 and blue when <0.5). The inward ownership becomes stronger from the first iteration and the BOWN computation reaches equilibrium quickly.

**Figure 5:**
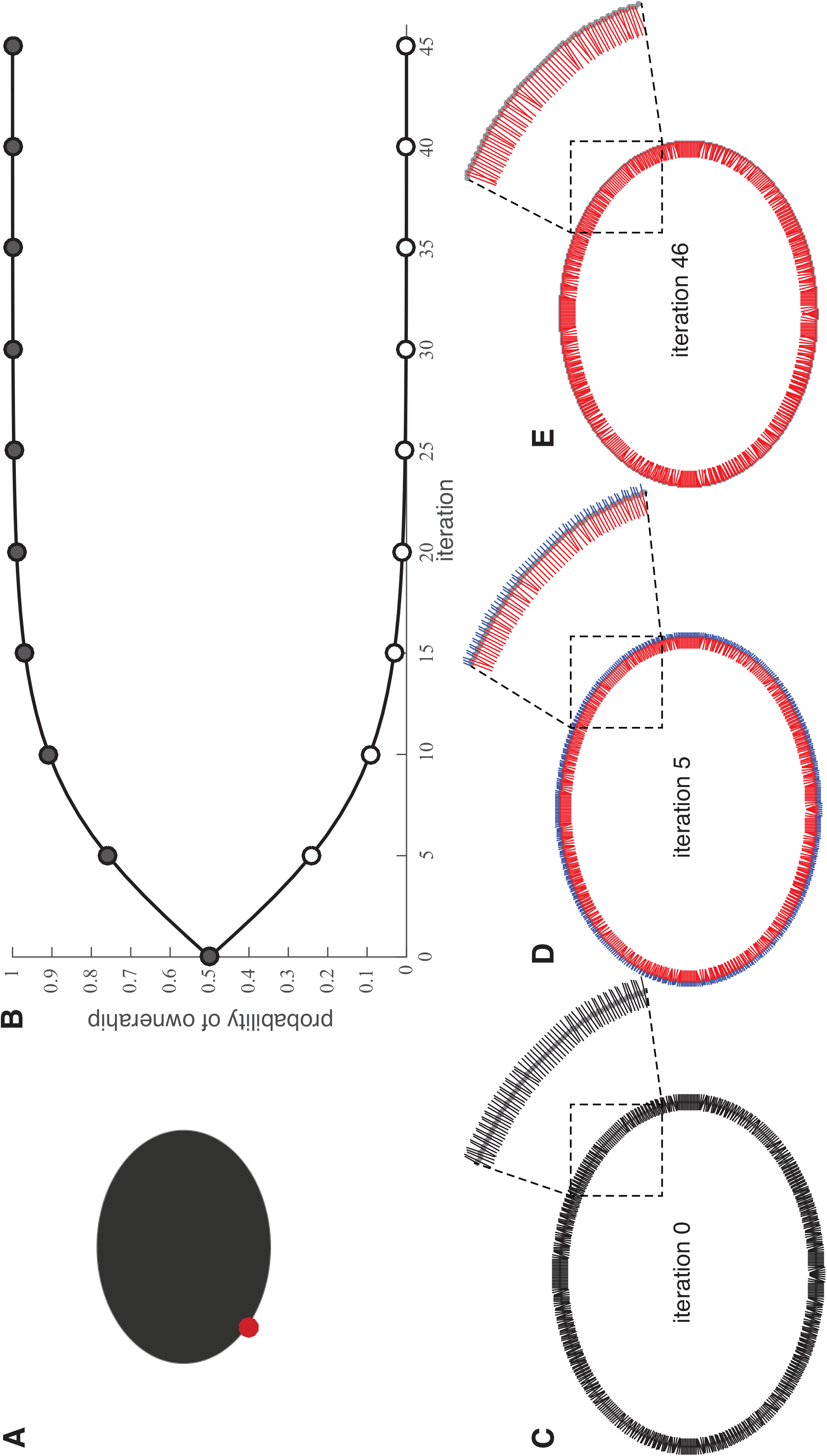
Response of the model to an oval shape. (A) Input image. The red-dot shows the location of the BOWN signal plotted in B. (B) The probability that the dark gray side and the white side of the border being the owner at the location of the red dot in (A) is plotted as a function of the model’s iterations. The circles in the plot are filled with the gray scale value of the owner-side. (C) The initial BOWN values (0.5 on both side) at iteration 0 are shown. Note that only a randomly sampled set of BOWN signals is plotted in the paper due to the high density of border nodes in input images, for visualization purposes. (D) and (E) As the iteration progresses a bias towards inside of the oval emerges, going through iteration 5 (D) until convergence at iteration 46 (E). Here, the red lines depict the winning side (BOWN signal value > 0.5), and blue the losing side (BOWN signal value < 0.5).

### Images with concavities

An oval is a compact shape, locally convex at any point along the contour. The results shown in Figure 5 are therefore not surprising with the convexity bias implemented in the algorithm (and are equivalent to the previous models’ responses). However, even if a surface has a large concave part, such as shown in Figure 6A, it is perceived as a figure. The BOWN signals for the inward ownership (black side) on the right-sided border (*B_2B_* and *B_3B_*) are inconsistent with each other, but consistent with the BOWN signals for the inward ownership on the left-sided convex border (*B_1B_*). The BOWN signals for the outward ownership (white side) on the right-sided border (e.g., *B_2W_* and *B_3W_*) are also consistent with each other. This causes a strong competition at the right-sided border. However, because the BOWN signals on the left side outnumber the BOWN signals on the right side (the border length on the left is longer than the one on the right), inward BOWN signals on the left border enhance the BOWN signals for inward BOWN on the right border. Hence, they become the winner of the competition eventually (Figure 6D). The time courses of the development of BOWNs at the point *P_1_* and *P_2_* in Figure 6D is shown in Figure 6B and C, respectively. The time course of the BOWN signals shows the slower development of inward ownership on the right (concave) side than the left (convex) side due to the competition with the outward ownership.

**Figure 6:**
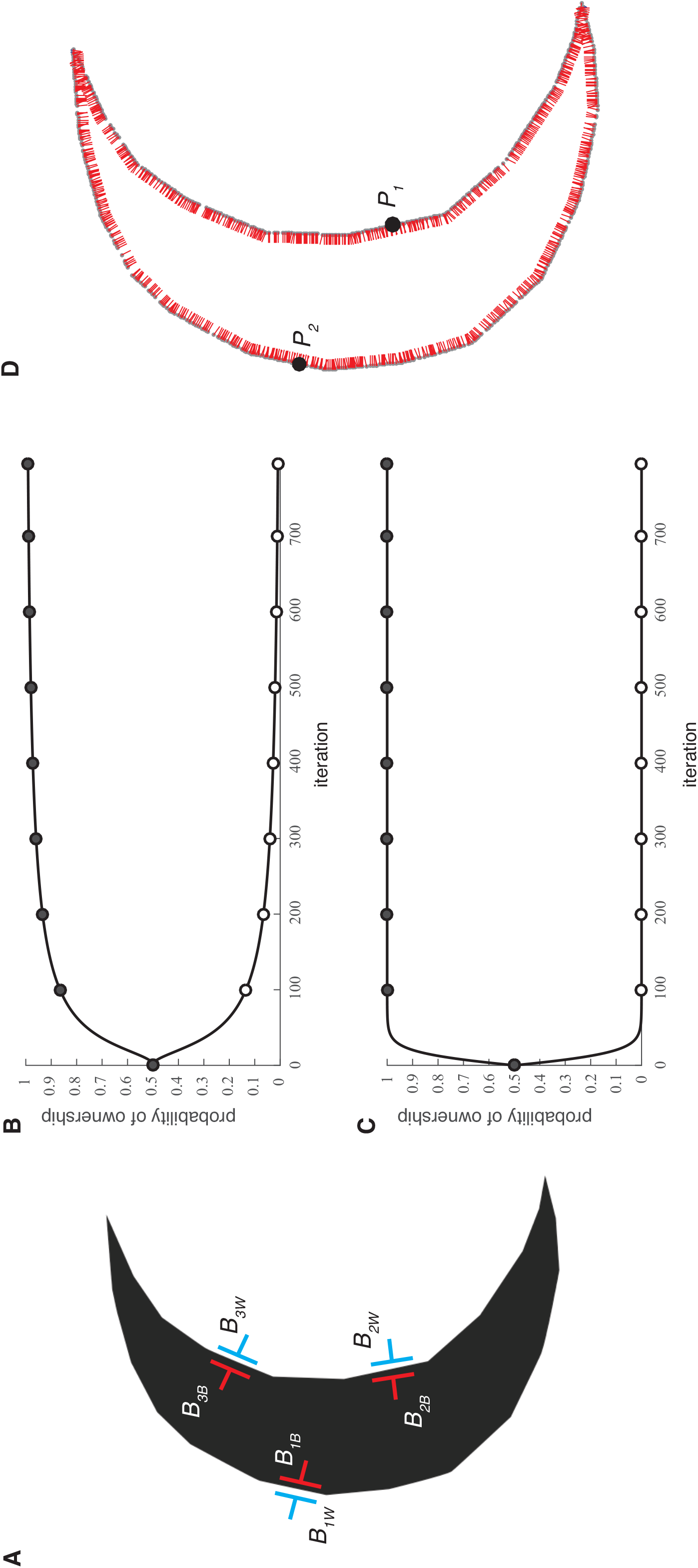
The response of the model to a crescent moon shape. (A) Input image with BOWN signals referred in the main text. (B) The probability of the competing BOWN signals at location *P_1_* in (D) as a function of the model’s iterations. (C) The probability of the competing BOWN signals at location *P_2_* in (D). (D) The model’s response at convergence.

Next, we will consider images shown in Figure 4, equivalent to Figure 2. In Figure 7A, the inner area of the image has a different color compared to the other areas equivalent to the image in Figure 2A. In contrast, in Figure 7D, the inner area has the same color as the background, equivalent to Figure 2B that likely evokes the “hole” perception. The BOWN signals at equilibrium are shown in Figure 7B and 7E, respectively. The time course of BOWN signals at the equivalent points on the inner borders in Figure 7B and 7E are plotted in Figure 7C and 7F, respectively. For Figure 7A, BOWN signals quickly reach equilibrium with the inward directions of the ownerships for both inner and outer borders ([suppl movie 3](https://www.dropbox.com/s/2vr4onuit203nzx/movie_suppl_3_donutInconsistent2_opt.gif?dl=0)), indicating that the black and grey areas are both figural. In contrast, the development of BOWNs at the inner border in Figure 7D shows a more complex time course. It starts with weak dominance of the inward ownership for several iterations and then is reversed to outward ownership ([suppl movie 4](https://www.dropbox.com/s/kpbznaha9p2y0vv/movie_suppl_4_donutConsistent2_opt.gif?dl=0)). It also takes a longer time to reach equilibrium than in Figure 7A. The final BOWN indicates that the central area is a hole and the black area is a donut-shaped figure. It is also important to note the difference of the BOWN values of the inner border at the equilibrium between Figure 7C and 7F. Although the outward BOWN signals in Figure 7D are clearly dominant, the final BOWN values do not reach 1.0 (as in Figure 7C). This indicates that the perception of the hole in Figure 7D is less decisive. In general, when an area is perceived as a hole the model often shows the initial inward dominance and later reversal to outward dominance. This takes more iterations to reach equilibrium, reflecting the gradually increasing influence of global interactions. These observations likely correspond to the general ambiguous perception of holes. As discussed in the introduction, the two images have exactly the same border maps, and hence, previous models without reflecting surface properties would not be able to distinguish the different perceptual organizations for the two images, indicating the strength of our approach.

**Figure 7:**
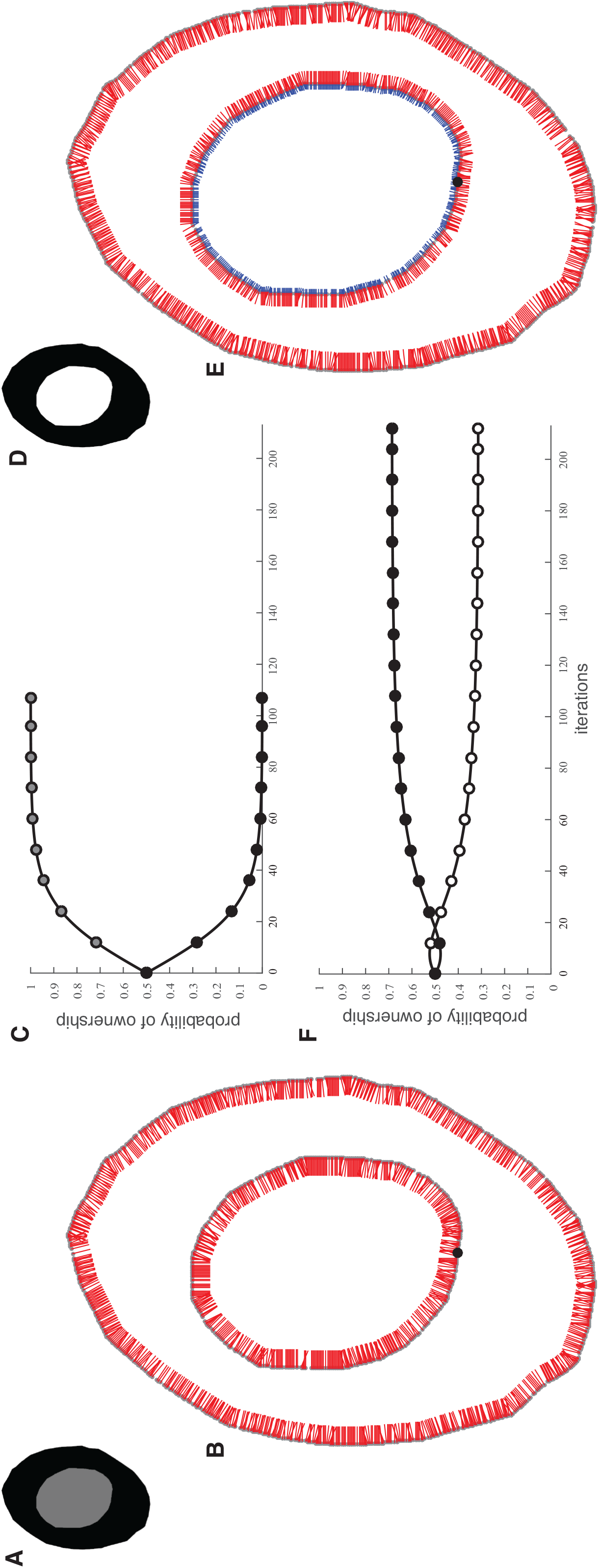
The response of the model to overlapping ovals (A) and a donut shape (D). (B) Model’s response at convergence for image in (A) showing ownership of the inner border by the gray internal oval area. (C) The time course of the competing BOWN signals of image in (A) at the location indicated by the black dot in (B). (E) Model’s response at convergence for image in (D) showing outward ownership of the inner border by the black donut area. (F) The time course of competing BOWN signals at the location indicated by the black dot in (E) (the same location as in (B)). Note the reversal of ownership from inward to outward direction at around iteration 20.

Next, we analyze the responses of the model to a less compact shape: an elongated curved shape with convex and concave parts (Figure 8A, convex and concave parts are indicated by red and blue arrows, respectively). Examining the response of the model to such a shape is essential, because, if a model with a long-range interaction has only a convexity bias algorithm, it would work for compact shapes, but not for elongated shapes with a mixture of convex and concave parts (such as this S-shape). In Figure 8B, the magnified part within the dotted square in Figure 8A is shown. Although the border on the right side is concave, the leftward BOWN signal (green) is consistent with the rightward BOWN signals on the border on the left side (orange) based on the convexity detection algorithm. Hence, they are enhanced. In Figure 8C, the response of the model is shown. Because the overall length of the convex border is longer than the one of the concave border, the inward BOWNs become the winner at all locations of the borders at equilibrium. Figures 8D and E show the time course of the development of BOWN signals for BOWN at *P_1_* on the concave part, and at *P_2_* on convex part in Figure 8C, respectively. The ownership of the concave border starts with the outward direction in the first few iterations, but dominance is quickly reversed and the inward ownership becomes the winner. A comparison between Figure 8D and Figure 8E shows that the ownership of inward BOWNs develops more slowly at the concave border. This difference in the time course of the development can also be observed in [suppl movie 5](https://www.dropbox.com/s/mb51fcjl56bjue0/movie_suppl_5_snake1_half_opt.gif?dl=0). The result is in clear contrast to the computation solely based on a convexity bias as discussed later where the responses of the model with and without the surface consistency constraint are compared.

**Figure 8:**
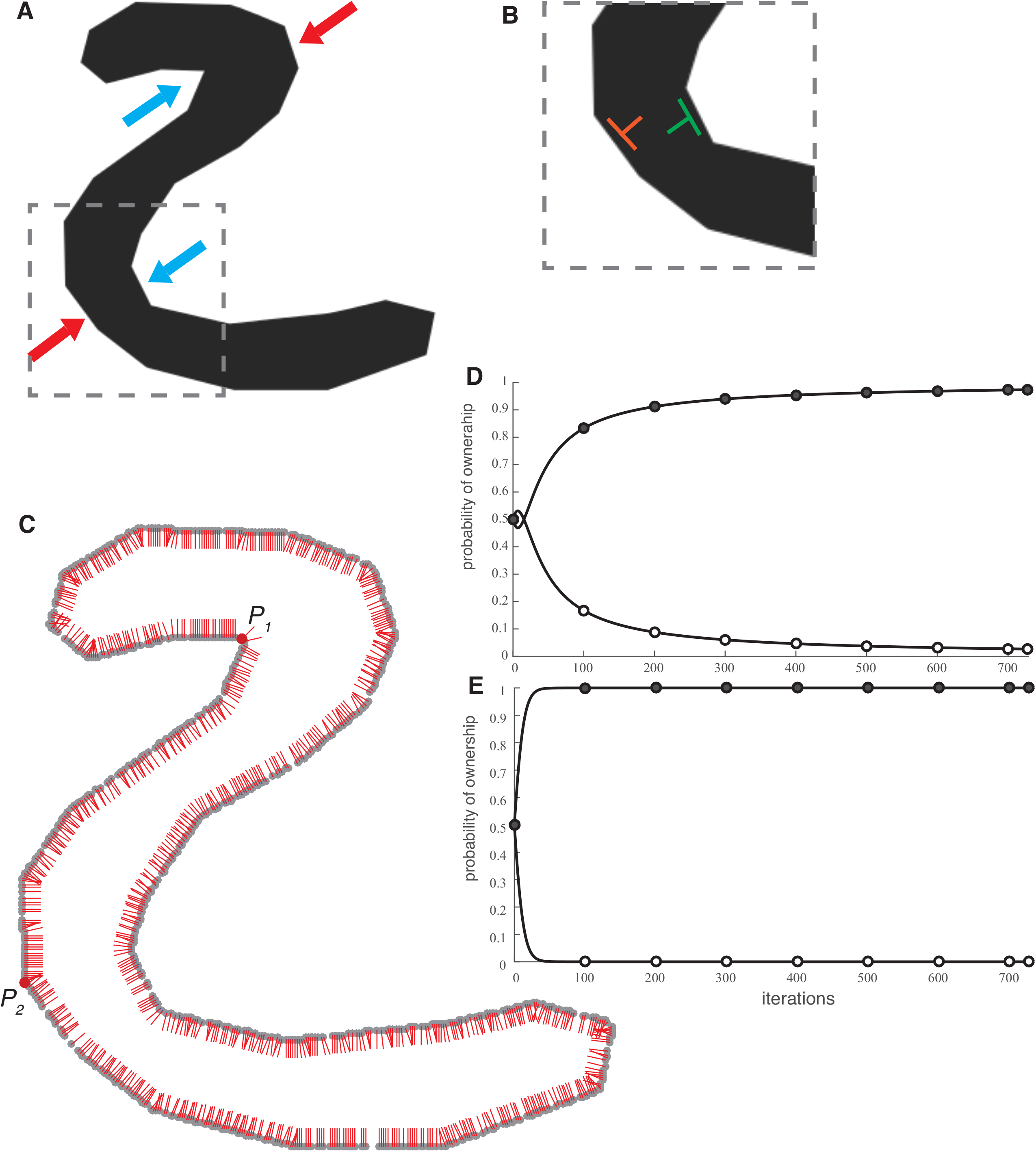
Model’s response for S-shape. (A) Input image. Red arrows: convex part. Blue arrows: concave part. (B) Zoomed-in of the dashed square area in (A). (C) Model’s response at convergence. (D) Time course of the probability of the black side and the white side being the owner at location *P_1_* in (C). (E) Time course of the probability of the black side and the white side being the owner at location *P_2_* in (C).

(See Supplemental Material Figures S2A-B for additional data with images that have many concavity parts.)

Furthermore, we investigated the effect of the enclosedness of the concave area (Figure 9A, B, and C). While the shape in Figure 9A has a wide opening, the other two images have a narrower opening, and Figure C has a nearly-closed concave area (similar to the images used in Kim & Feldman, 2009). The surface areas of the black area in the three images are kept constant. The time course of the BOWN signals for the white-sided ownership at the center of the concave border in Figure 9A (red dot) and at the equivalent positions in Figure 9B and 9C are shown in Figure 9D. The plots in Figure 9E, data from iteration 0 to 20, shows that the ownership of the center of the concave border was stronger on the white side at first (the BOWN value larger than 0.5 in the plot indicates ownership by the white side) and only after a certain number of iterations, was it reversed. Although in all cases the black area became the winner at equilibrium, the ownership by the black side is the weakest in the image in Figure 9C (the lower value in the plot in Figure 9D indicates the stronger ownership by the black side). The result suggests that the likelihood that the observer reports the concave area as a figure is higher with a higher degree of closure (Figure 9C>9B>9A), similar to the results by Kim & Feldman (2009).

**Figure 9:**
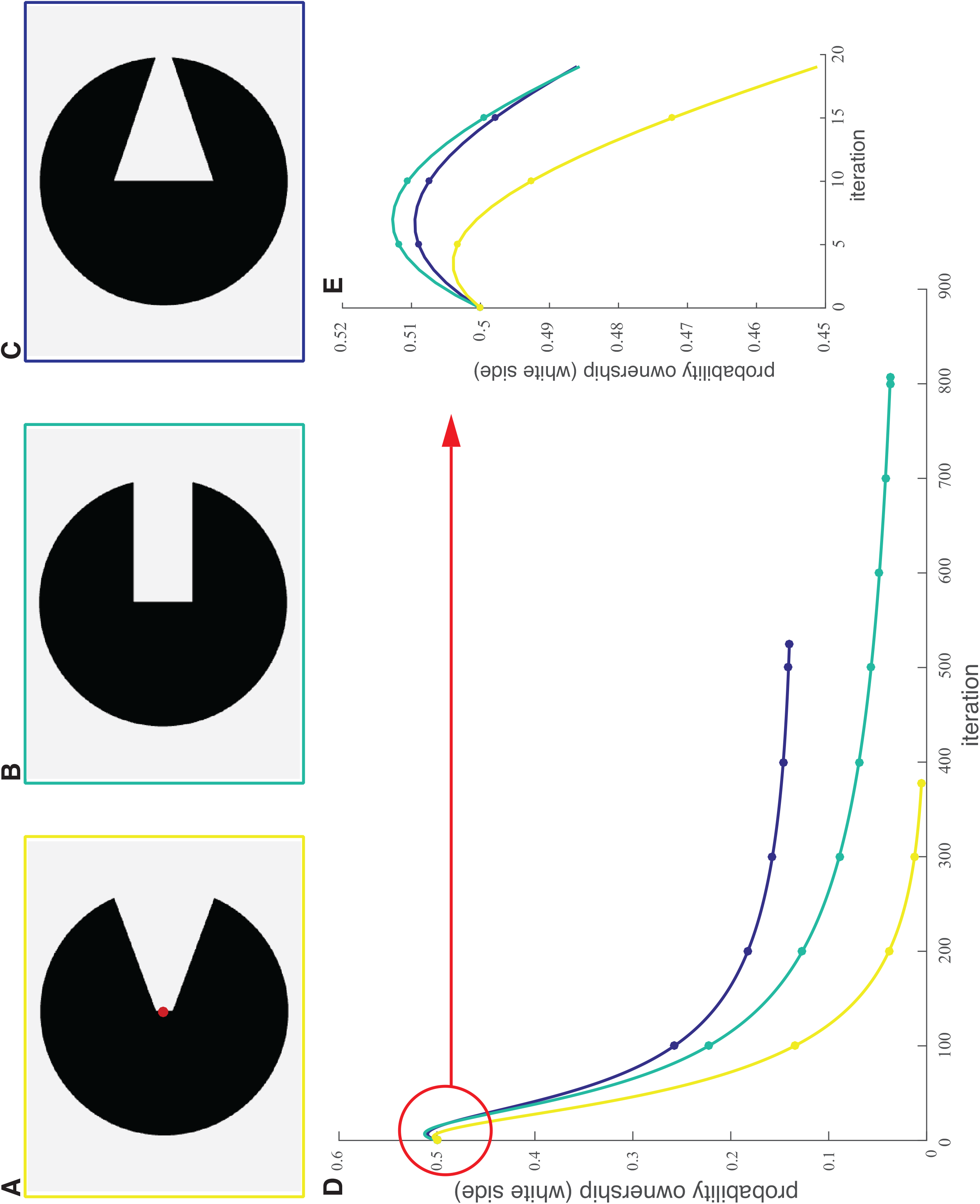
Model’s responses for C-shapes with different concavities at the opening part. (A-C) Input images. (D) Time course of the probability that the light grey area is the owner at the location indicated by the red dot in A. The colors of the plots correspond to the color of the square frames in (A-C). (E) Magnified plot of the part indicated by the red circle in (D).

### Images with occlusions

Next, we describe responses of the model to images in which occlusions of surfaces are perceived. Figure 10A shows a stereotypical image that gives rise to a perception of occlusion. However, when the vertical oval is moved downward (as shown in Figure 10B), the border between the two areas is no longer shared, and two separate objects are perceived (see Bushnell et al., 2011, who used similar images and showed differential neural responses in V4). The model’s response shows that the border shared between the two areas in Figure 10A is owned by the vertical oval, i.e. it occludes the gray surface (Figure 10C). The gray surface is the owner of the part of the border that is shared with the white background. In response to Figure 10B, inward BOWN signals became the winners all along the two borders (Figure 10D), indicating that these two areas are separate figures.

**Figure 10:**
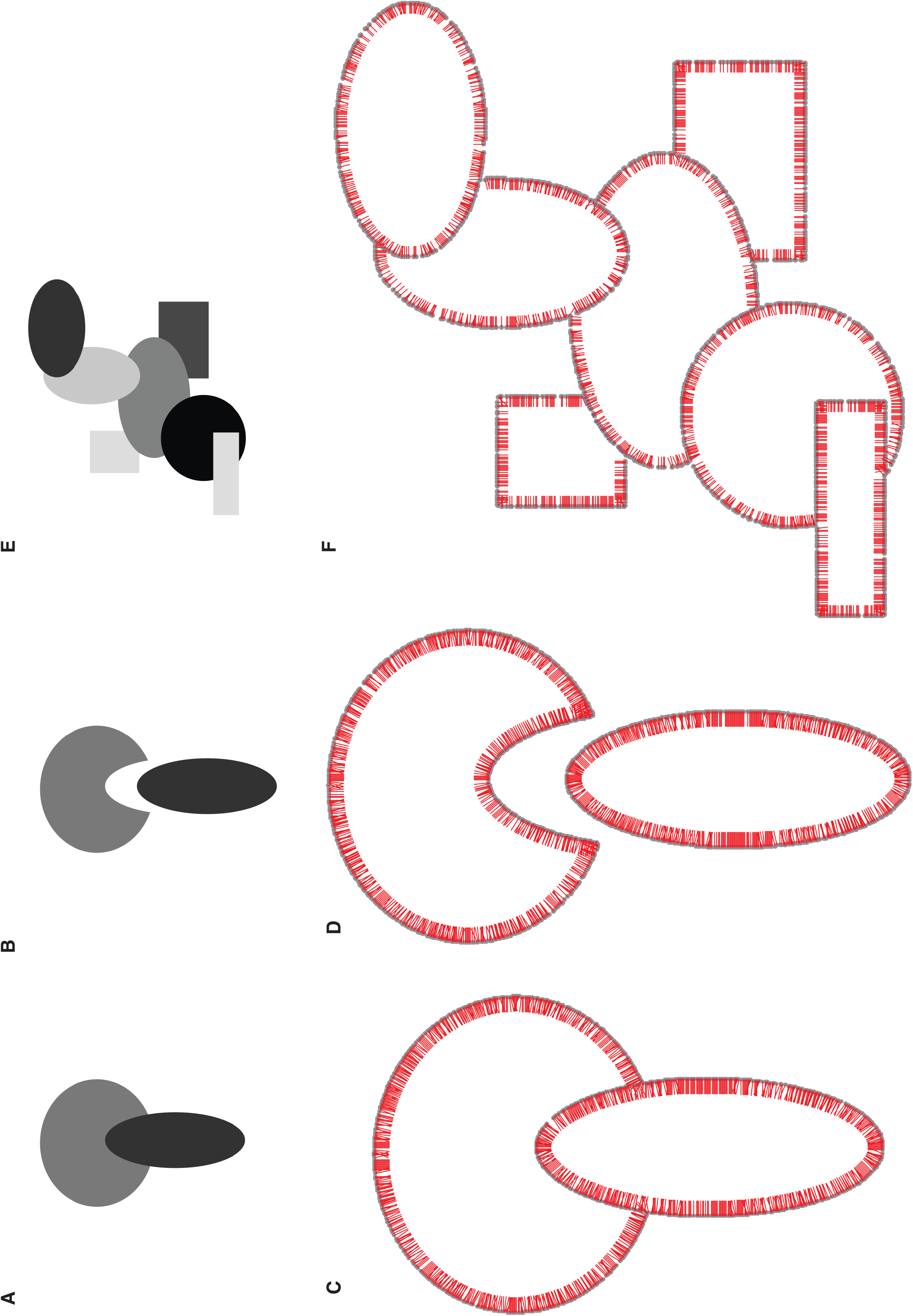
Model’s responses to images with perceived occlusion. (A) Input image perceived with occlusion. (B) Modified image in (A) to avoid the perception of occlusion. (C-D) Model’s responses at convergence to images in (A and B), respectively. (E) Input image that is perceived as overlapping multiple surfaces. (F) Model’ response to the image (E).

Even when multiple overlapping surfaces are present (as shown in Figure 10E), human vision can detect occlusion with ease. However, if a model is based solely on a convexity bias, it would have difficulties reproducing this perception. To resolve this problem, many previous models added T-junctions as an additional cue for occlusion (Calderero & Caselles, 2013; Dimiccoli & Salembier, 2009; Thielscher & Neumann, 2008; Zaidi, Spehar, & Shy, 1997, but see Calderero & Caselles, 2013; Dimiccoli, 2015). Due to the surface consistency constraint in our model, selective and specific (but not accidental) interactions between consistent BOWN signals are made, and our model produces responses with inward directions of BOWN signals in the individual surfaces without the addition of T-junction detectors (Figure 10F). When the surface consistency condition is removed from the model, the result shows incoherent BOWN signals within the individual surfaces as discussed later.

As shown earlier, the model is capable of reproducing the perception of a hole. What if an occluded object is perceived through the hole? Figure 11A is an example of such a condition. In one of the two oval areas in the black surface, the part of the contour of the occluded dark gray surface is perceived. The other oval area in the black surface is perceived as a surface on top of the black surface. The response of the model is shown in Figure 11B. The BOWN signals with inward directions at the contours of the dark gray, black, and light gray surfaces (blue arrows) are the winners. On the other hand, outward BOWN signals are the winners at the border of the oval hole area (green arrow) indicating that it is a hole. At the border segment within the hole, winning BOWN signals are the ones with upward direction, indicating that it is a part of the contour of the dark gray object. The reason why they become the winner is because these BOWN signals are consistent with the BOWN signals of the dark gray surface (purple BOWN signals) due to both the convexity relationship and the surface consistency, and hence, selective interactions occur between them. Without the surface consistency constraint, many accidental interactions would occur between the existing BOWN signals in this image.

**Figure 11:**
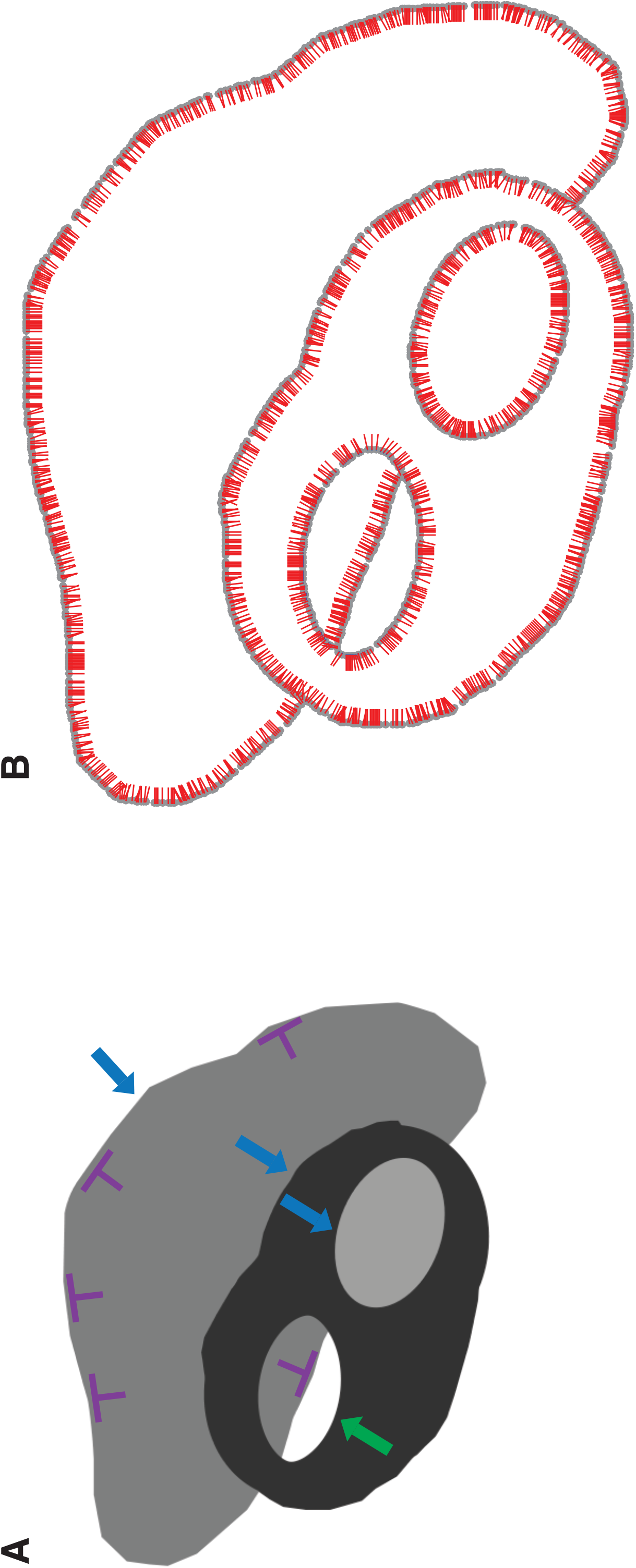
(A) An image where the left side oval area is perceived as a hole with a part of the contour of the dark gray surface is seen. See main text for the BOWN signals at the locations indicated by the blue, and green arrows. BOWN of the border segment within the hole area is perceived to belong to the dark gray side. (B) Model’s response at convergence. The oval area on the right is owned by the inward BOWN signals while the oval area on the left is owned by the outward BOWN signals. The border segment within the hole is owned by the dark gray side because of the consistency of the BOWN signals with the other inward BOWN signals of the dark gray area (purple BOWN signals in (A)).

Images in Figure 12A and B evoke a further complex perceptual organization. In Figure 12A, white circular areas are perceived as holes because the background is also white, except where one of the circular areas in gray (bottom right) is perceived as a surface on top of the black surface, because the gray color is not consistent with the rest of the image. In Figure 12B (modified from Kellman, 2003), multiple holes in the black surface and an occluded dark gray surface are perceived. Furthermore, the light gray circular area in the middle of the bottom row is perceived as a surface on top of the black surface due to the difference of the color from the rest. As shown in Figure 12C and 12D, the model gives responses that correspond to the perceptions: outward ownership for the hole areas, and inward ownership for the figural areas, including the fractions of contours of the occluded surface (see [suppl movie 13](https://www.dropbox.com/s/4606knx1avrn86n/movie_suppl_13_holeMesh_opt.gif?dl=0) and [suppl movie 14](https://www.dropbox.com/s/r2osikg7486r1pl/movie_suppl_14_holeMeshObject5_opt.gif?dl=0)).

**Figure 12:**
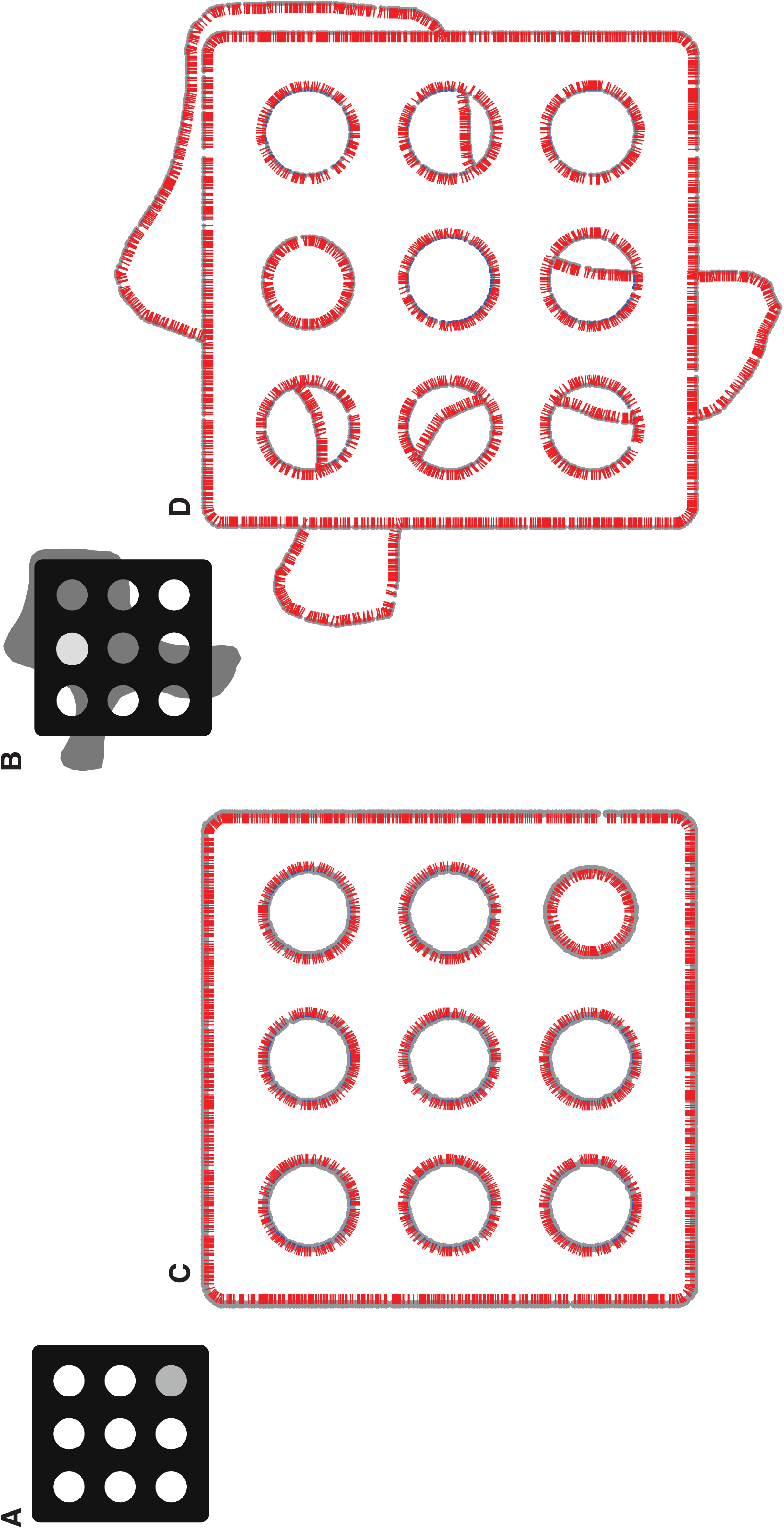
(A) An image that is perceived with multiple holes in the circular areas, except the right bottom circular area where its color is different from the rest and hence perceived as a surface on top. (B) An image as in (A) except a dark gray surface is perceived as occluded by the black surface with multiple hole areas. The top middle circular area has a different color from the rest and hence perceived as a surface on top. (C) and (D) Model’s responses at convergence of input images (A) and (B) respectively. Note the ownerships of border segments within the hole areas by the dark gray sides.

### Relatability

Curvilinear alignment of border segments at two intersections corresponding to the perception of occlusion was defined as “relatability” by Kellman and Shipley (Kellman & Shipley, 1991). In Figure 13A, a black rectangle is perceived to occlude a gray rectangle. BOWN signals of border segments that intersect with the occluder, *B_1_* and *B_2_*, are colinearly aligned (i.e. they are relatable). In Figure 13B and 13C, the image is modified such that the relatability of *B_1_* and *B_2_* is violated. However, the BOWN signals of inward directions at both border segments satisfy the convexity conditions with many other BOWN signals at other border segments (e.g., *B_3_*, *B_4_*). Furthermore, many of them satisfy the convexity condition with *both B_1_* and *B_2_* at the same time (orange lines). This means that, for example, *B_1_* enhances *B_3_* which, in turn, enhances *B_2_* and vice versa through the iterations. Therefore, *B_1_* and *B_2_* influence each other indirectly. As the result, the model shows robust responses to images with occlusions even when the borders at intersections are not aligned (Figure 13E, F).

**Figure 13:**
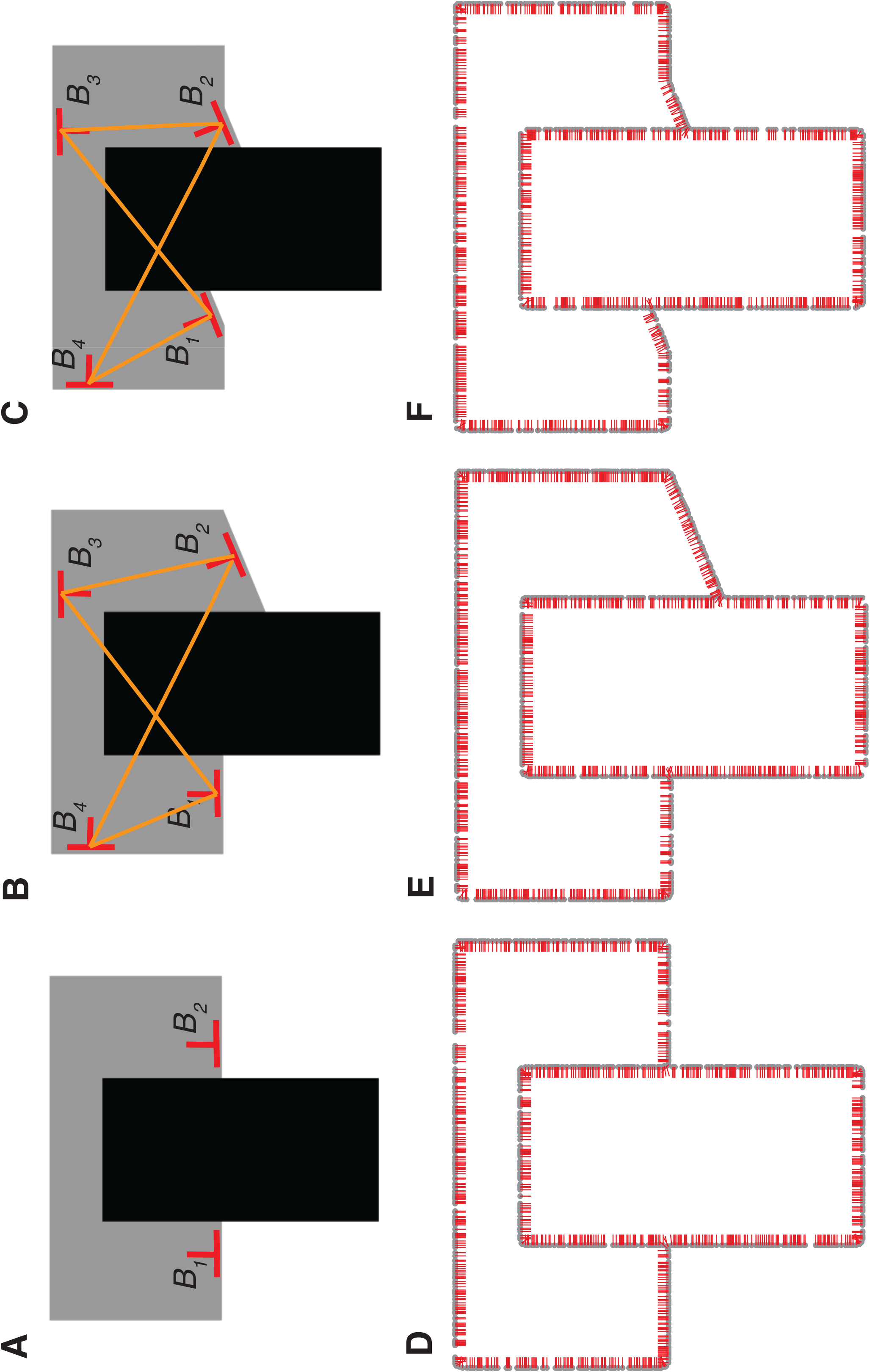
Relatability of border segments and BOWN computation. (A) An image that is perceived with occlusion. BOWN signals on border segments on the left and on the right of the occluder (black rectangle), *B_1_* and *B_2_*, are aligned colinearly. The border segments satisfy the relatability relationship (Kellman & Shipley, 1991). (B-C) By changing the orientation of the border segment on the right (B) or on both sides (C), they do not satisfy the relatability relationship and *B_1_* and *B_2_*, are no longer aligned. However, they interact with other inward BOWN signals of the gray area such as *B_3_* and *B_4_*. As the result, *B_1_* and *B_2_* are enhanced. (D-F) Model’s responses at convergence of input images (A-C).

In the image in Figure 14A, a part of the gray surface is perceived through the hole of the black ring-shaped surface. In Figure 14B∼I, the part within the hole area is rotated clockwise at 22.5° increment. In this way, the BOWN signal of the border within the hole area, *B_1_*, becomes non- relatable to the BOWN signals of the two border segments at the intersections outside of the ring-shaped surface, *B_2_* and *B_3_*. In Figure 14J-R, the responses of the model to these images are shown. As the rotation angle increases, the ownership of the border is at first on the gray side. However, at 67.5°, this manipulation causes some BOWN signals of the inner circular border to become inward along the border (Figure 14D, orange arrow). This is due to the fact that the outward BOWN signals at the inner circular border (e.g. *B_4_*) are no longer consistent with the inward BOWN signals of the outer circular border (e.g. *B_5_*) due to the inconsistency of the un- owned side colors. The inconsistent BOWN signals along the circular border increase as the rotation angle increases (Figure 14E-14I). At 180°, inward BOWN signals become dominant at the entire inner circular border. The ownership of the straight border at the center becomes ambiguous (equal amount of ownerships for both directions, orange arrow) because they are not consistent with any other BOWN signals except themselves. It is possible that these results correspond to the ambiguity of the perception of the circular area caused by this manipulation. The results presented in this section indicate that BOWN signals at the location of intersection with the occluder are enhanced in our model, even when they do not have the “relatable” partner-BOWN signals. This is because they have other BOWN signals that satisfy the constraints of convexity and surface consistency. In other words, our model is tolerant, to a certain degree, to misalignment of border segments which may correspond to human perception as experienced in Figure 13 and 14 (see Supplemental Material Figures S2C-F for additional data).

**Figure 14:**
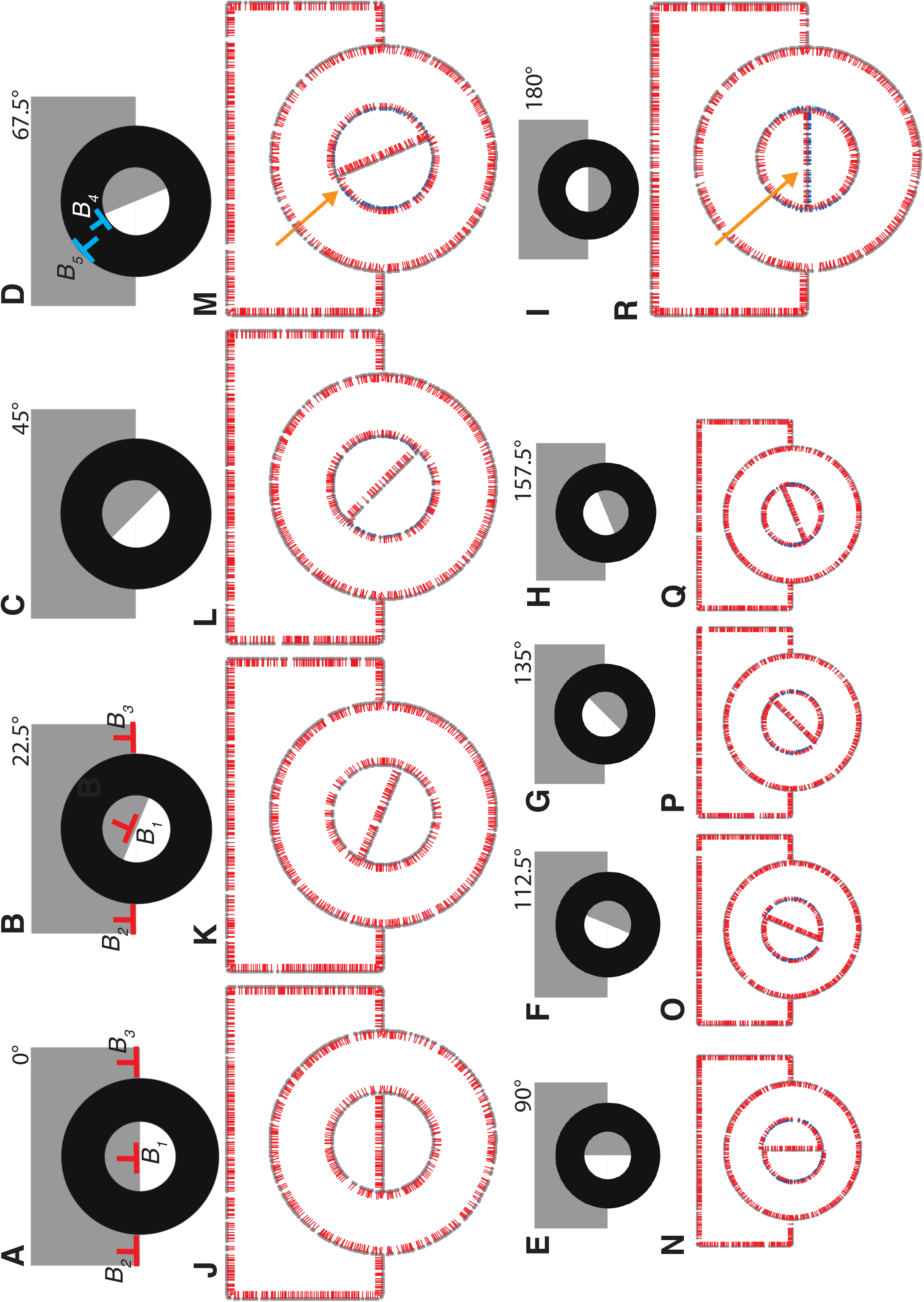
Misalignment of border segment within a hole area and BOWN computation. (A) Top: an image where a hole in the black circular area and a border segment of a gray rectangle within the hole are perceived. *B_1_*, *B_2_* and *B_3_* are all colinearly aligned. (B-I) Top: images as in (A) except the gray area within the hole being rotated in clockwise by 22.5° increment. Due to the modification, the border segment within the hole is no longer “relatable” with the ones one the left and on the right. (J-R) Model’s responses to the images in (A-I). As the rotation progresses, there are more inconsistencies between the outward BOWN signals of the inner circular border (*B_4_* in (D)) and the inward BOWN signals of the outer circular border (*B_5_*). As the result, BOWN signals become inconsistent along the inner circular border (orange arrow in M). When the rotation is 180° (I), the inward BOWN signals become dominant at all locations of the inner circular border. The ownership of the straight border segment within the circle is ambiguous and its ownership is divided by both sides.

### Role of long-range interactions: surface consistency constraint and space constant

As shown above, the implementation of the large-scale surface consistency constraint gives robust responses. To further analyze the effect of this implementation, the responses of the model with and without the surface consistency constraint are compared.

In the image with a perceived hole (Figure 4B, shown again in Figure 15A), if the surface consistency constraint is not implemented, the border of the hole area becomes owned by the inward direction. This is because “accidental” interactions occur between unrelated BOWN signals that have convexity relationship. Consider a BOWN signal, *B_2i_* (red), on the border of the hole area. This signal is in competition with the other BOWN signal, *B_2o_* (blue). With the surface consistency constraint, *B_2i_* is consistent with the other BOWN signals of the same inner border with inward directions (e.g. *B_1i_*, red), while *B_2o_* is consistent with the BOWN signals on the outer border with inward directions (e.g. *B_3i_*, blue). When the latter interactions supersede the former, the outward BOWN signals of the inner border such as *B_2o_* become the winner. However, without the surface consistency constraint, *B_2i_* can interact with the other BOWN signals (such as inward BOWN signals located on the part of the outer border above *B_2i_* (e.g. *B_4i_*, blue signal on top). As the results, *B_2i_* becomes stronger than *B_2o_* as shown in Figure 15C.

**Figure 15:**
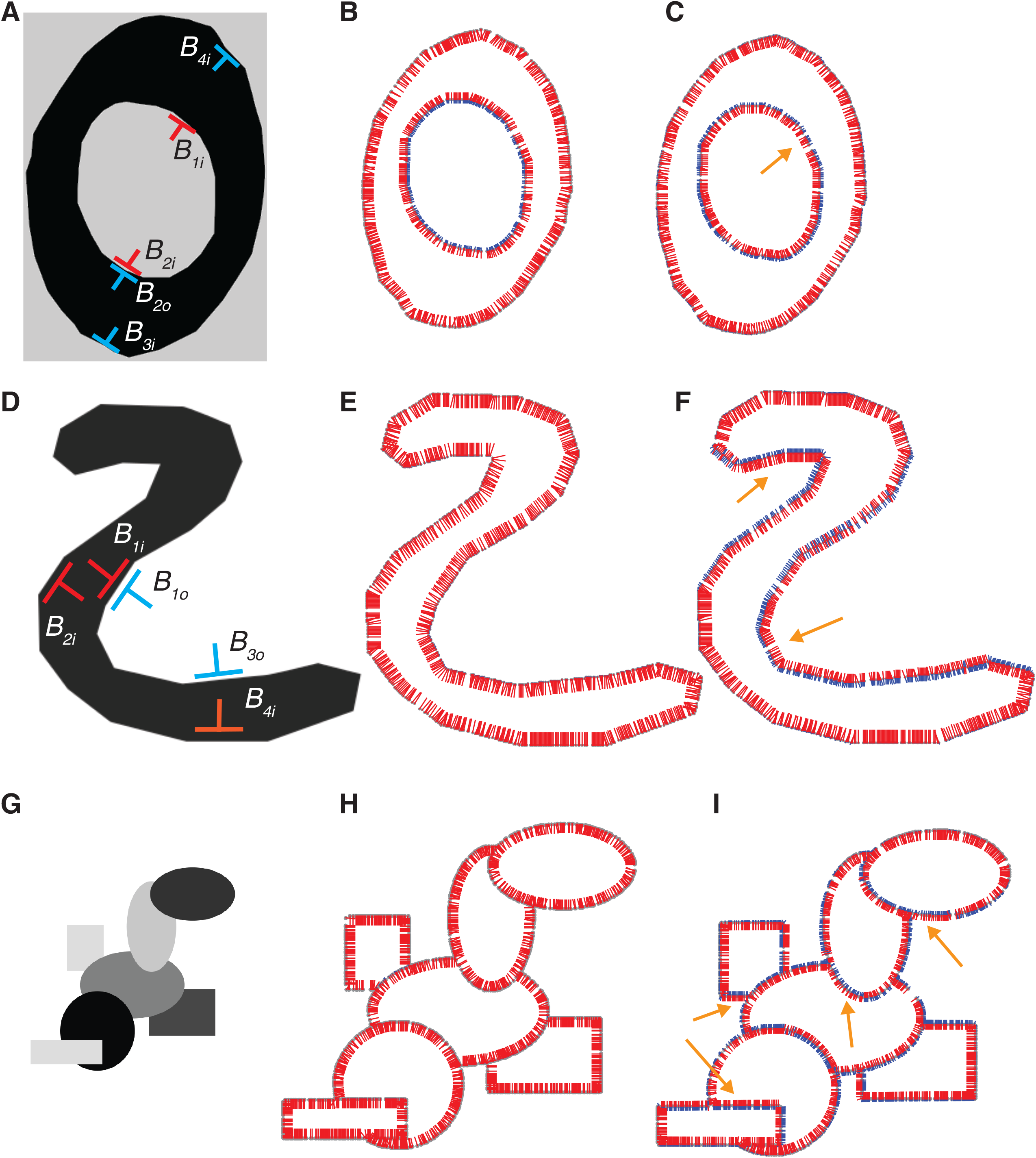
Influence of surface consistency condition of the model’s responses. Left column: input images to the model. Middle Column: output of the model at convergence with surface consistency constraint. Right column: output of the model at convergence without surface consistency constraint. Orange arrows indicate incorrect BOWNs due to the lack of surface consistency detection.

Consider next, the response of the model to the S-shape used earlier (Figure 15D). Without the surface consistency constraint, there are also accidental interactions of BOWN signals. In the competition of BOWN at a location on a concave contour, the BOWN signals (e.g. *B_1i_* and *B_1o_* (Figure 15D)), the interactions between *B_1i_* (red), and the other inward BOWN signals (e.g. *B_2i_* red), supersede the interactions between *B_1o_* (blue) and the other outward BOWN signals on the concave contour (e.g. *B_3o_*, blue), if the surface consistency constraint is implemented. However, without the surface consistency constraint, *B_1o_* starts to have accidental interactions with other BOWN signals such as the one on the bottom contour, *B_4i_* (orange). The surface color of the owned side is white in *B_1o_* and black in *B_4i_*, and vice versa for the colors of the un- owned side. Hence, the interactions between the two signals did not occur when the surface consistency constraint was implemented. Without it, however, these signals interact. As the result, the strength of *B_1o_* becomes stronger than the competing *B_1i_* (orange arrows in Figure 15F).

In the image with multiple overlapping surfaces (Figure 15G), accidental interactions occur in many locations if the surface consistency constraint is not implemented. As the result, there are BOWN signals indicating outward direction of ownership (Figure 15I orange arrows). These examples clearly indicate the necessity of implementing the surface consistency constraint in the long-range interactions of BOWN signals.

Next, we examined the effect of the space constant value, *σ* (eq. 3), to the BOWN computation. If the value is large, it makes the long-range interactions stronger, while if the value is small, the interactions are limited to a local effect. This is illustrated in Figure 16 with the S-shape (Figure 16A). In Figure 16C, D and E, results of the computations with the space constant values of 20 pixels, 200 pixels (standard parameter of the model), and 1000 pixels are shown, respectively. When the space constant is smaller, the strong interactions between neighboring outward BOWN signals on the concave part of the contour (Figure 16B, blue BOWN signals) make the outward BOWN signals competitive with the inward BOWN signals. This is because the inward BOWN signals on the same contour (pink BOWN signals) do not interact with each other directly because they do not satisfy the convexity condition and only have long distance interactions with the signals on the other side (red BOWN signals). As a result, outward BOWNs become more dominant in some parts (orange arrows in Figure 16C). The result indicates that it is essential to set the *σ* value high, relative to the size of the input image so that long-range interactions can be effective. In doing so, the model is able to reflect global configurations. On the contrary, when the space constant is large enough, it does not influence the results. Specifically, a larger space constant merely causes more iterations until convergence (Figure 16E).

**Figure 16:**
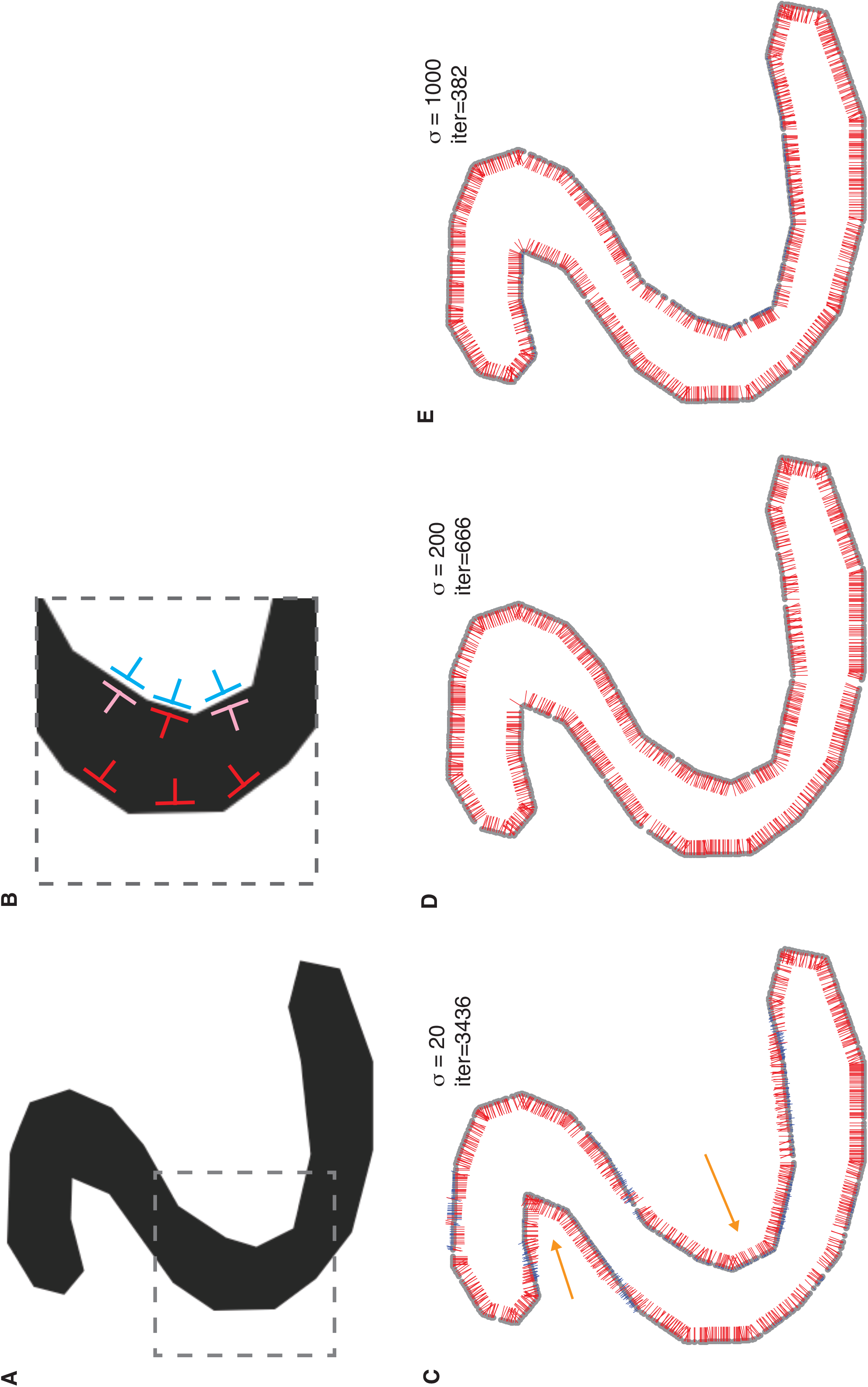
Influence of space constant parameter, *σ*. (A) Input image to the model (same as Figure 8). (B) A magnified part from the square in (A) with exemplar BOWN signals discussed in the main text. (C) Model’s response at convergence when *σ* = 20. (D) Model’s response at convergence when *σ* = 200 (the standard parameter used in this paper). (E) Model’s response at convergence when *σ* = 1000.

We also analyzed the condition where there are separate surfaces with same color in which, in principle, the BOWN signals of separate surfaces interact in our model. In Figure 17A, two surfaces with same color are placed next to each other. The incorrect BOWN signal pair (blue), the leftward BOWN signal on the left-sided border of the circle (*B_3_*) and the rightward BOWN signal on the right-sided border of the rectangle (*B_4_*), are consistent with each other. However, their competitors (the correct BOWN signals (*B_1_* and *B_2_*)), have larger numbers of BOWN signals that they are consistent with (red BOWN signals, e.g. *B_5_* and *B_6_*). As a result, BOWN signals with inward directions of the two surfaces become the winner (Figure 17B). Furthermore, there are also accidental interactions between correct BOWN signals in two separate surfaces (incorrect interactions between correct BOWN signals). The inward BOWN signals of the right-sided border of the circle (e.g. *B_5_*) and the inward BOWN signals of the left-sided border of the rectangle (e.g. *B_6_*) are consistent. As described in the introduction, separate BO cells interact without the continuation of a border between the border elements they respond to (Zhang & von der Heydt, 2010). Accordingly, our model lets BOWN signals interact without the constraint of continuity. Hence, the accidental interactions, such as between *B_5_* and *B_6_*, can occur if there are multiple separate surfaces with same color. Note that these accidental interactions are incorrect interactions between correct BOWN signals in separate surfaces and have the following characteristics: (1) even though they interact, their effects are in the correct directions; (2) because their distance is longer than the correct interactions within the individual surfaces, those closer would be the more dominant factor for interactions.

**Figure 17:**
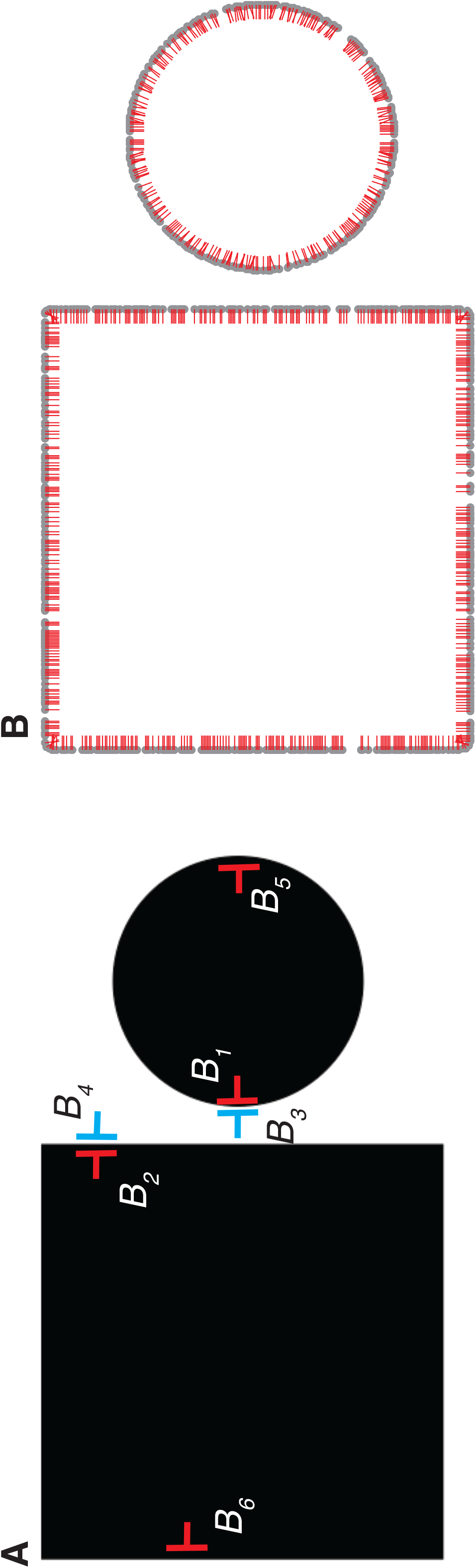
Response of the model to an image with two separate areas with same color. (A) Input image and example BOWN signals discussed in the main text. (B) The response of the model at convergence.

Hence, when there are accidental interactions between incorrect BOWN signals, the effects are overwhelmed by the interactions between correct BOWN signals within each individual surface. When there are accidental interactions between correct BOWN signals from separate surfaces, the outcome does not, in effect, disturb the correct BOWN computation. However, it is quite intriguing to see how the neural system deals with the accidental interactions between correct BOWN signals described above. It is left to future research to investigate if there are non-linear dynamic grouping mechanisms which enhance correct interactions between correct BOWN signals to further facilitate BOWN computations.

### An example with an inconsistent response

As described above, the implementation of the global surface consistency constraint, along with the convexity constraint, gives robust responses. However, we also encountered cases where the model is not able to reproduce human perception. A representative example is shown in Figure 18A: the image is perceived as a part of a gray oval covered by a black crescent moon-like shape. The response of the model shows that the border shared by the black and gray areas is owned by the gray side (Figure 18B, orange arrow) which does not correspond to the perception of occlusion. The colors of the two sides of the border are dark gray and black.

**Figure 18:**
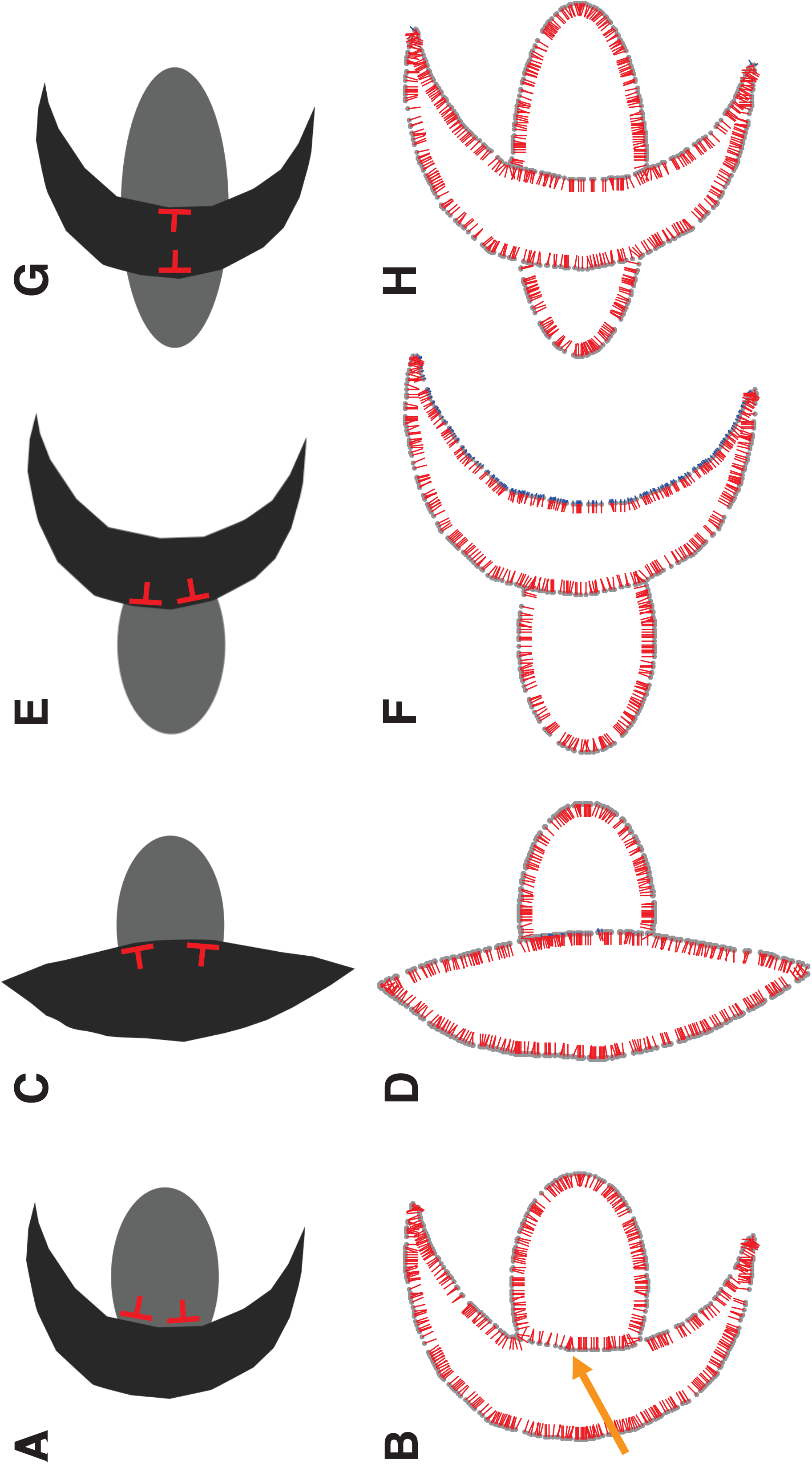
A problem of the approach implemented in the standard DISC2 model. (A) An image where a gray oval is perceived to be occluded by a black crescent moon. The perceived occlusion occurs at the concave part of the crescent moon. (B) The response of the model showed that the border segment between the black and gray areas is owned by the side of the gray area, the result that does not match the perception of occlusion. (C, E) Input images modified from (A). The gray oval is perceived to be occluded by a convex shape (C) or at the convex part of the crescent moon (E). (D, F) Model’s responses to the images in (C and E), respectively. Because the BOWN signals at the border between the gray and the black areas is now owned by the black side because the inward BOWN signals (red) satisfy the convexity relationship. (G) An image modified from (A) by elongating the gray oval so that a part of it is seen on the other side of the crescent moon. There are two border segments between the black area and the two gray areas. There are interactions between the inward BOWN signals of black area on the two border segments (red). (F) The response of the model to the image in (E). Because of the interaction, the inward BOWN signals for the black area become the winner.

Because of this combination of colors, BOWN signals on this part of the border are not consistent with any other BOWN signals in the rest of the image, due to the surface consistency constraint. Hence, the interactions occur only between BOWN signals within the border segment. As the result of the convexity bias, the BOWN signals on the dark gray side become the winner. In contrast, if the crescent moon shape is replaced with a convex shape (Figure 18C) or if the gray oval is occluded on the convex side of the crescent moon (Figure 18E), the model’s responses show that the BOWN signals on the black side are the winner (Figure 18D and 18F, respectively). Furthermore, if the oval in Figure 18A is elongated horizontally and a part of it appears on the other side of the crescent moon (as in Figure 18G), the model’s response indicates the occlusion of the oval by the crescent moon shape (Figure 18H). In this case, because there are two border segments shared by the black and gray areas, the BOWN signals on the right-sided border, and the ones on the left-sided border (red signals) now have consistent surface colors. Hence these BOWN signals with inward directions enhance each other. As the results in Figures 18D-F show, the inward BOWNs become the winner in these examples. The problem in Figure 18A originates from the fact that the BOWN signals at the border segment do not interact with the rest of the BOWN signals outside the segment, and hence, global configurations do not influence this part of the BOWN computation. This condition, the abutment of an area to a convex part of a border of another area, can appear frequently in images with natural and/or complex shapes (see Figure 21). We will show a solution for this problem by the “augmented model” in the following text.

### Model convergence

We analyzed the extent to which the model’s convergence depends on the input images. Figure 19 shows the number of iterations it took to reach equilibrium for all images used in this paper. This plot gives a general idea of what causes the model to take more iterative processes: complexity (the right-most two images with holes and occlusions) and ambiguity (due to the existence of concave contours (S-shape)) causes more competitions between opposing BOWN signals and hence, it takes longer for the model to equilibrate^3^.

**Figure 19:**
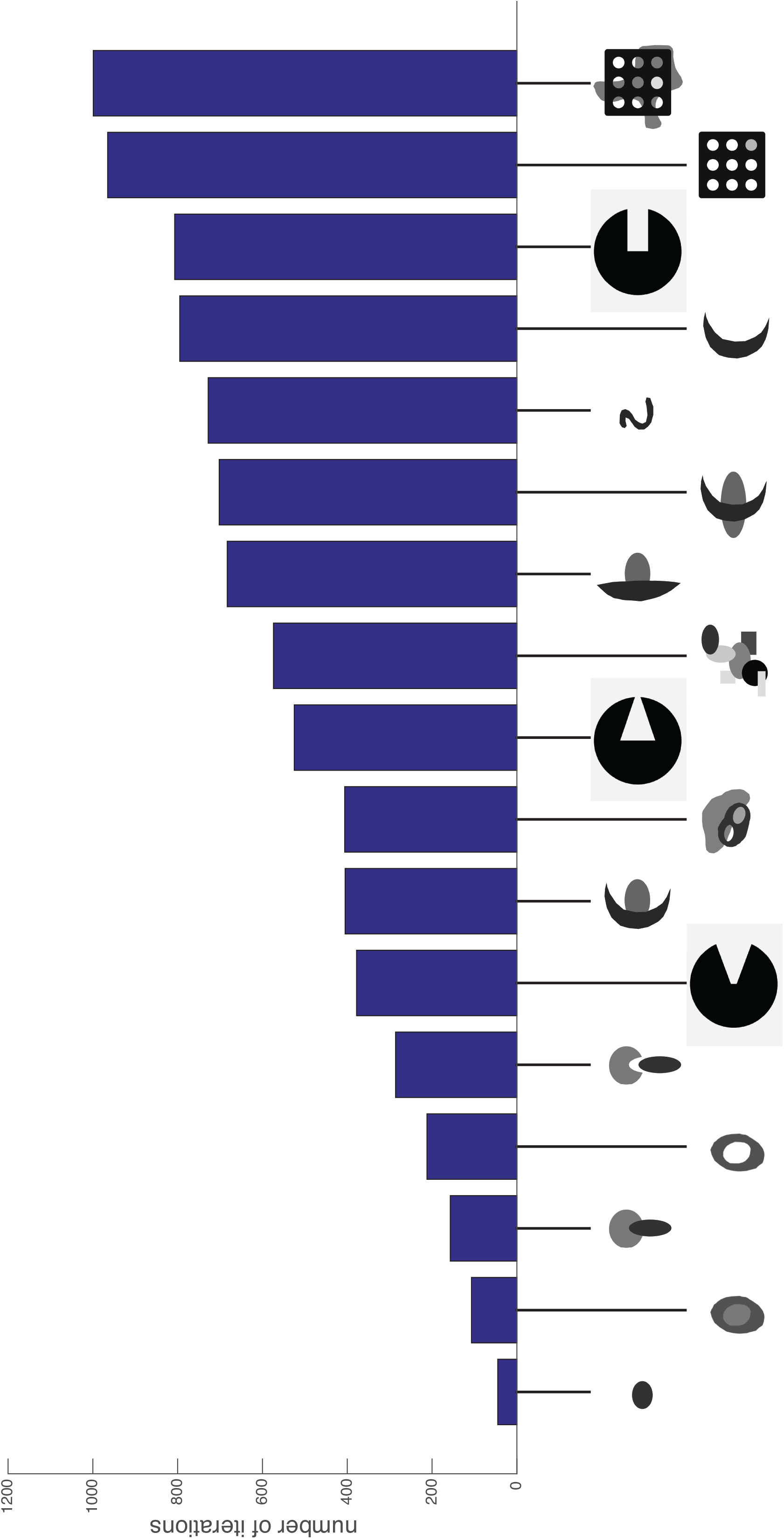
Summary of the number of iterations until convergence for all images used in this paper.

### Augmented model

The problem of the inconsistent responses described above can be solved by constructing the two-layered “augmented DISC2” model. In addition to the BOWN computation based on the global interactions implemented in the standard DISC2, a local, curvilinearity-based interaction between neighboring BOWN signals is implemented. In this computation, the nearby BOWN signals enhance each other if (1) they are curvilinearly aligned (with same owner side) and (2) they have the same surface color on the owner side. The image from Figure 18A and BOWN signals at around the T-junction of the image are shown in Figure 20A and 20B, respectively. In the global BOWN computation, *B_1_* and *B_2_* are not considered to be consistent due to the inconsistency of the surface properties on the un-owned side. Consequently, in the standard DISC2, only *B_4_* and *B_6_* among the shown BOWN signals, are directly enhanced. However, *B_1_* and *B_2_* (red) are curvilinearly aligned and have a consistent surface color on the owner-side. Hence, they are enhanced in the local BOWN computation in the augmented DISC2. Note that the colinearly aligned BOWN signals on the other side, *B_3_* and *B_4_* (blue) are not enhanced due to the inconsistency of the owner side surface colors. The enhancement between *B_1_* and *B_2_* means that mutual enhancement goes beyond the location of the T-junctions (green circle) where the color on the un-owned side changes. *B_1_* is further enhanced by the long-range interactions with *B_7_* (in Figure 20A) in the global BOWN computation. This in turn enhances the local interaction of *B_1_* with *B_2_*. The response of the augmented DISC2 to the image is shown in Figure 20C. The border at the concave side of the crescent moon shape is now owned by the interior of the shape. The problem of the standard DISC2 occurs when occlusion is perceived at a concave contour (well-represented by Figure 18A). This problem is now solved in the augmented DISC2 by combining the global computation with the local curvilinearity-based computation.

**Figure 20:**
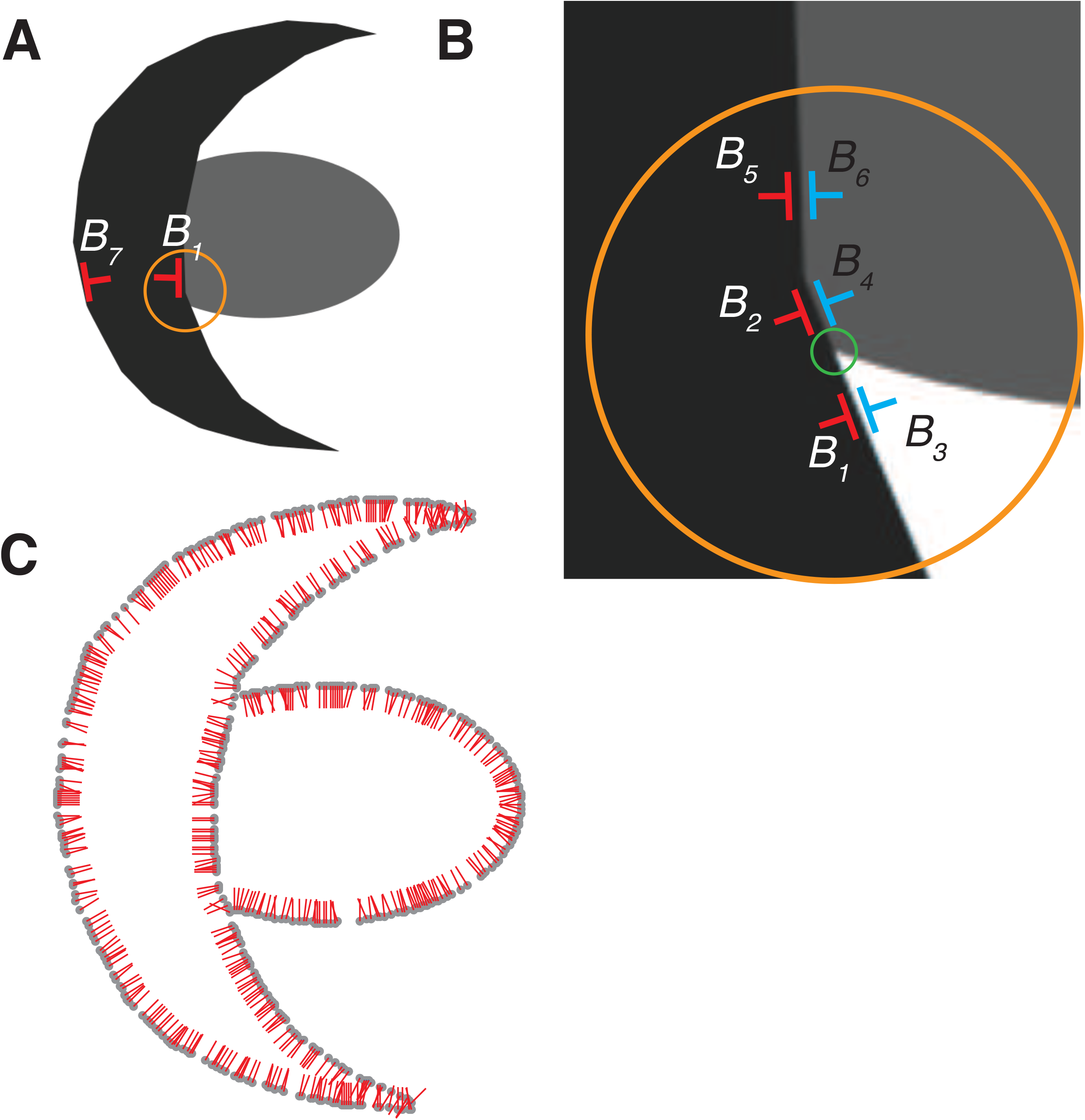
Approach of the augmented DISC2 model and its response. (A) The image shown in Figure 18A which the standard DISC2 model failed to reproduce the perception of occlusion. (A) The relationships of BOWN signals at around the T-junction indicated by the orange circle in A. *B_1_* and *B_2_* are curvilinearly aligned and the owner side color is same. Hence, in the local BOWN computation in the augmented DISC2, the two BOWN signals enhance each other. In addition, *B_1_* is enhanced by the global BOWN computation by the interaction with the BOWN signals on the other side of the black area, e.g. *B_7_* in A. (C) The response of the augmented DISC2. The border segment between the black and the gray area is owned by the black side. Compare this with the result shown in Figure 18B.

In Figure 21A, an image modified from a well-known image by Bregman (Bregman, 1981) is shown. In this image, there are many parts where the occluder side has a concave shape, as shown in the magnified images (blue circles). In the magnified images, the locations of intersections of the perceived occluded contours are marked with orange circles. The responses of the augmented DISC2 to this image is shown in Figure 21B, showing correct BOWN signals along all borders. This is in a strong contrast with the response of the standard DISC2 to the same image shown in Figure 21C. In Figure 21D and E, another variation of the Bregman image and the response of the model are shown, respectively. Although the response shows correct BOWNs for the majority of locations, in some locations, BOWNs are shown in the opposite direction, or shown without clear dominance on either side (orange circles). This indicates that there are some cases where even the augmented DISC2 does not show fully- coherent BOWNs.

**Figure 21:**
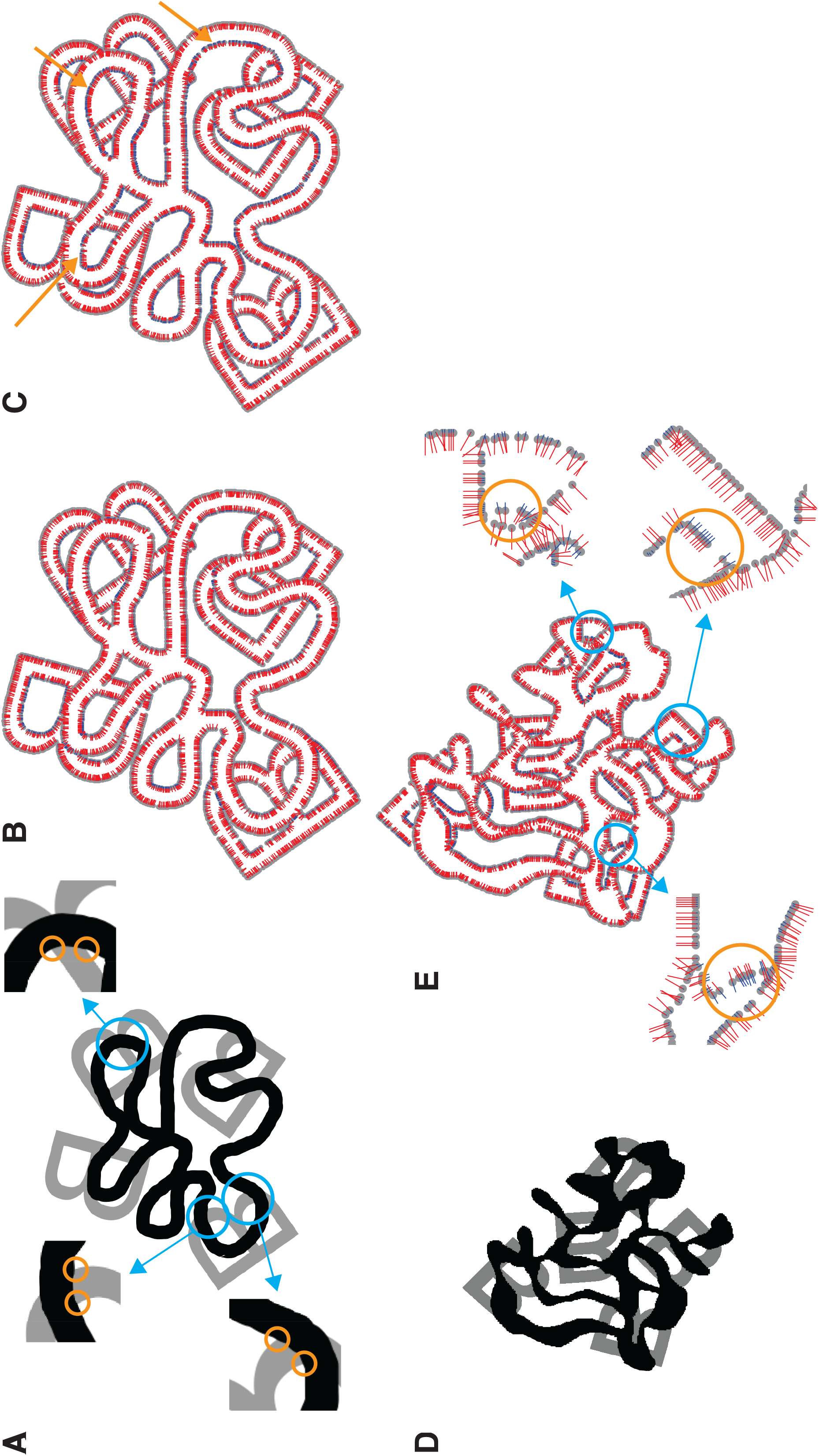
Responses of augmented DISC2 to modified Bregman images (Bregman, 1981). (A) An example of a modified Bregman image where perceived occlusions occur at concave parts of the black occluder (blue circles). The parts are magnified and the T-junctions at the occlusions are indicated by orange circles in insets. (B) The response of the augmented DISC2 model. (C) The response of the standard DISC2 model (orange arrows indicate inconsistent BOWNs). (D) Another example of modified Bregman image. (E) The response of the Augmented DISC2 model to the image in (D). Although at majority part of the borders the response showed correct ownerships, there are responses at some locations (blue circles) that were not coherent with the rest (orange circles in the magnified parts of the blue circles).

The responses of the standard and augmented DICS2 to all the images used in this paper are available at (https://github.com/vickyf/disc2).

## Discussion

The model reported here was developed to compute BOWN signals that underlie figure-ground organization. As discussed in the introduction, the BOWN signal reflects global configurations of input images. To reproduce this property, the model takes the following approaches, 1. long-range interactions: all combinations of all BOWN signals in the image are considered for interactions, no matter how separated they are in distance, and 2. rules of interactions: certain constraints of interactions should be applied to specifically enhance globally-coherent signals. For this purpose, three constraints of interactions between BOWN signals were implemented: the pair of BOWN signals has to have a convexity relationship, have consistent surface properties, and the interaction strength decreases exponentially by the distance between the pair. The approach with surface consistency detection was inspired by the “augmented model” of Zhaoping (Zhaoping, 2005) where she implemented an algorithm that detected a consistency of contrast polarities between two BOWN signals. This model allowed only these consistent BOWN signals to interact. This implementation reflected the fact that a large portion of BOWN sensitive neurons (75%) are also contrast-sensitive and, only when the preferred contrast is present, the neurons showed BOWN tuning (Zhou et al., 2000). In this paper, we examined the robustness of such an approach. If a model starts solely from the border maps of images, only geometry-based constraints (such as the convexity constraint) are applicable, and hence, is bound to fail in the reproduction of figure-ground organization of many images (e.g. Figure 7). By adding the capability to import images and compute border nodes and their orientations, we were able to examine BOWNs of complex shapes (Zhaoping’s model and the previous DISC model had a limit of detecting only vertical and horizontal orientations of border segments). In contrast to many previous models without surface consistency detection, our model showed unprecedented robust responses. The robustness of this approach is clearly observed in its responses to the images with complex depth orders that include areas perceived as holes, occlusions, and multiple overlapping surfaces (Figure 7, 10E, 11, and 12). The results strongly suggest that surface consistency detection, in conjunction with the convexity bias, is essential in perceptual organization (Bertamini, 2006; Bertamini & Hulleman, 2006; Yin et al., 1996, 2000), and that the contrast-sensitive BOWN neurons and their long-range interactions may underlie the processes for surface consistency detection.

However, the problem of this approach was illustrated by counter examples: when an occlusion is perceived at the concave part of an occluder, the standard model is not capable of indicating the ownership of the border by the occluder (Figure 18B). The solution to this problem was given by an implementation of a two-layered model with local and global computations in the augmented DISC2 model. In the local computation of the model, no matter whether a pair of BOWN signals forms a convex or concave contour, they are enhanced as far as they are curvilinearly aligned, share same owner side, and have the same surface property on the owner side. This extends the interactions of neighboring BOWN signals beyond the T-junction at the location of occlusion. They are further enhanced by the long-range interactions in the global BOWN computation, which, in turn, enhances the local interaction. The dynamic interactions between the local and global computation may be linked to the function of feedback signals from a higher-level to help the coherent integration of lower-level signals. “Synergistic” interactions between feedback projection and lateral connections in a context-sensitive manner in the visual cortex have been investigated (Bullier, 2001; Gilbert & Li, 2013; Hupé et al., 1998; Lamme, 1995; Liang et al., 2017; Roelfsema et al., 2002; Roelfsema, 2006; Zhou et al., 2000). It is possible that dynamic interactions between lower-level and higher-level computations are involved in BOWN computation, where the enhancement of signals occurs only when the lower-level signals are coherent with the higher-level signals in terms of the owner side of borders, as implemented in the augmented DISC2.

Next, we will discuss how the model is able to show robust responses despite not having specific junction detectors. T-junction detection has been a common approach to reproduce the perception of occlusion (Calderero & Caselles, 2013; Dimiccoli & Salembier, 2009; Thielscher & Neumann, 2008; Zaidi et al., 1997 to name a few). However, there are two issues to be considered. First, there is no clear evidence that a T-junction detection mechanism exists in the visual system. Second, a behavioral study indicated that occlusion is perceived only when the context surrounding a T-junction is presented, but not when a T-junction is presented within a small window (McDermott, 2004).

To address the first issue, note that the T-junction consists of three abutting surfaces: the abutment of two surfaces creates the “stem”, and their abutment with another perpendicular surface creates the “top” of T-junction. Therefore, neurons tuned to a T-shape (for example Brincat & Connor, 2004) are not a T-junction detector. One possible candidate of a T-junction detector is an end-stopped cell. It has been argued that end-stopped cells play a key role in figure-ground organization (Craft et al., 2007; Heitger et al., 1992; Heitger & Heydt, 1993). Furthermore, it has been suggested that contrast-sensitive end-stopped cells in V1 (Yazdanbakhsh & Livingstone, 2006) are involved in signaling at a T-junction. However, for the end-stopped cell to work as a T-junction detector, it has to detect the three-surfaces configuration as described above, and it is not clear how these neurons respond to such a configuration. For the second issue, based on the lack of perception of occlusion by the T-junction itself, McDermott suggested the possibility that T-junctions might not be detected prior to, and thus not used for, scene interpretation. Furthermore, Tse reported various images where a part of a 3D shape is perceived to continue behind another object without explicit T-junctions (Tse, 1999). Note that DISC2 model reflects the configuration of a T-junction without specific T-junction detectors, due to the consistency of the surface property of occluded surface. Furthermore, in the augmented DISC2, the collinearity detection is combined with the surface consistency detection on the owner side. This algorithm gives the area above the “top” part of T-junction an advantage in achieving the ownership of the “top” border.

The question is, what features of images should be detected in a model for figure-ground organization. The issue can be formalized as follows: Consider four algorithms: convexity detection, C, surface consistency detection, S, collinearity detection, L, and T-junction detection, T. As we reported here, the standard DISC2 has C and S, while the augmented DISC2 has C, S and L. They show robust responses including images with holes. Conventional models with C and T are not capable of reproducing the perception of a hole in Figure 6. Then, the question is whether the addition of T is necessary or redundant. On one hand, it is significant that our model shows robust responses and that T-junction *per se* does not evoke the perception of occlusion. On the other hand, as evolution progresses in a way that the biological system takes redundant strategies, it is possible that a yet-to-be-found T-junction detection mechanism is somewhere in the neural system and works in a complementary manner (but not in a deterministic manner) for figure-ground perception. The results shown in Figure 21E, BOWNs in some parts do not correspond to human perception, may suggest such a complementary system is present and the visual system can respond appropriately to complex sensory information from nature. This is a vital point not only for perception research, but also for applications in computer vision. The issue of whether a T-junction detector exists in our neural system, (if so,) why the detection itself does not evoke a perception of occlusion, and how it contributes to perceptual organization, should be investigated thoroughly in future research.

In a model for BOWN computation, developed to reproduce illusory surface perception (DISC model, Kogo et al., 2010), it was hypothesized that the perception of illusory surface (Kanizsa square) is mediated by the emergence of BOWN signals at the gaps between the image elements (inducers), and that the emergent BOWN signals constitute so-called illusory contour completion, or “modal” completion, (Michotte et al., 1964). In other words, the illusory-contour-sensitive neurons, IC cells, (von der Heydt et al., 1984) are, or are driven by, BOWN sensitive cells (BO cells, Zhou et al., 2000)). Because the BOWN computation reflects global configurations by long-range interactions, the differential perceptions of the illusory Kanizsa square image and its non-illusory variation were explained. As discussed later, it has been suggested that the BOWN computation is performed by dynamic interactions between BO cells, and feedback signals reflecting higher-level computation (mediated by “grouping cells”, Craft et al., 2007; Zhou et al., 2000). This has an important implication in the activation of neural signals at the location of illusory contour. Assuming the hypothesis above, where IC cells are identified as BO cells, the following sequence of neural processes may occur when the illusory surface image is presented. At first, the BO cells receive no sensory input at the gaps between the image elements. However, global computation at a higher level detects the configuration of an (illusory) surface. It then sends top-down signals to activate BO cells at the lower level, that are coherent with the existence of the surface. Consequently, the formerly silent BO cells at the gaps of the image elements now become activated. This emergent activation of silent BO cells corresponds to the perception of the illusory surface and its contours.

How feedback signals can activate silent BO cells precisely at the location of illusory contour without activating other non-relevant BO cells is a fundamentally important question. In this regard, it should be noted that the augmented DISC2 implemented the interaction between the local and global computations. In other words, the model required an existence of signals based on the local properties. Only then the global computation is able to properly affect individual BOWN signals. This implies the possibility that the activation of silent BO cells emerges as the results of non-linear dynamic interactions between lower and higher-level signals. For example, it is possible that subthreshold depolarization of the membrane potential by local signals is combined with additional depolarization by feedback signals, triggering non-linear dynamics of neural responses mediated by the NMDA receptor and dendritic activity (Larkum, 2013; Larkum et al., 1999; Self et al., 2012; Wagatsuma et al., 2016).

Another type of completion phenomenon, “amodal completion” (Michotte et al., 1964), is perceived in the images used in the section for relatability in this paper. The output of the model indicates that BOWN signals of the border segments of the occluded surface at the intersection with the occluder (e.g. *B_1_* and *B_2_* in Figure 14A and *B_1_*, *B_2_* and *B_3_* in Figure 15A) are “grouped”, in the sense that they are considered to be consistent and enhance the interactions between them. We take the position that, in the case of amodal completion, the above-mentioned activation of silent BO cells by feedback signals in modal completion does not occur due to the stoppage of the extension of the colinear border segment. Hence, the emergent activity of BO cells behind the occluder does not occur. This corresponds to the fact that we do not perceive modal completion of the contours at the location (i.e. we do not “see” the contours). An intriguing example called “quasimodal completion” (Kellman, 2003; Kellman et al., 1998) represents simultaneous occurrence of modal and amodal completions in one image. Again, we predict that emergent activity of BO cells occurs at the location of modal completion, but not at the location of amodal completion due to the stoppage of, in this case, the illusory contours at the location of intersection of the contour with occluder. Underlying mechanisms of emergent neural activity, as the result of non-linear interactions between lower-level signals reflecting local properties and higher-level signals reflecting global properties, is fundamentally important in understanding the brain mechanisms which process sensory information. This question should be elucidated in future research.

Another important issue in BOWN computation is whether BOWN signals should be qualitative or quantitative. In the former case, BOWN signals only indicate the owner side categorically. In the latter case, BOWN signals indicate not only the side of ownership, but also the quantitative depth difference between the owned side and the un-owned side. This issue is closely related to the perception of 3D surfaces (volume perception). For example, in images where a volume of an object with 3D curvature of a surface is perceived, a gradual change of depth along the rim of the volume can be perceived (Tse, 1999). Another example is when a trapezoid area is perceived to be a slanted rectangle where the depth difference, with background changes along the border, is perceived if it is implied by, for example, the gradient of texture density. To reflect the gradual change of the depth along the contour, the BOWN signals should change strength accordingly, if this depth change is meant to be represented by BOWN signals. Hence, an important question is whether this quantitative property of depth perception is expressed in the activities of BO cells, or if it is added by another neural mechanism at another level. This aspect of BO cells has not been elucidated in neurophysiological reports.^4^ Our current model normalizes two competing BOWN signals at each border node. Hence, if anything, it may indicate “the strength of ownership” by the balance between the two competing signals. In this scheme, the different equilibrium values for different figures may only reflect the ambiguity of the ownership. On the contrary, the example of slant clearly indicates that depth may change quantitatively without the change of ambiguity of the ownerships. This may suggest that higher-level computations, such as the computation of 3D profile of surfaces, is done on top of the ownership computation to constitute quantitative depth perception. It is important to investigate how the visual system represents the quantitative profile of depth based on pictorial cues, how metric and orderly depth signals are integrated in the visual system, and how BOWN computations are involved (or not) in these processes. These questions need to be thoroughly investigated in future research.

There are few additional functions that may be implemented to improve the model. First, luminance values on a single surface may vary due to the gradient of illuminations in natural images. The model would detect this difference of luminance, and process it as an inconsistency of surface properties. To compute figure-ground organization of natural images (Hu et al., 2019), first the well-known inverse optics problem has to be solved so that the computation reflects only intrinsic properties of the surface (reflectance) but not the extrinsic component (illumination). Second, segmentation by texture is not implemented in the model. However, the segmentation is only at the first stage of computation in the model, and the rest is the same after segmentation. Implementation of algorithms of texture detection and segmentation from computer vision research would make the model even more robust.

Finally, it is important to ask how the global interactions algorithmically implemented in our model are realized in a real-life neural system. Craft et al. (Craft et al., 2007) suggested a neural mechanism in which signals from BO cells are integrated by a “grouping cell” (G-cell) with a ring-shaped receptive field (see also Hu et al., 2019; Wagatsuma et al., 2016). In the model, G-cells collect signals from BO cells that (1) are located within the ring area and (2) have a preferred owner side in the inward direction of the ring. Then, the G-cell sends feedback signals to the same BO cells to enhance their activities. In this way, the G-cell gives a bias to the lower-level competition between BO cells in the inward direction of the ring. This results in a convexity bias in their overall responses. This neural mechanism is one likely candidate of realizing the long-range interactions implemented in the model. It is also important to point out that there are contrast non-sensitive BO cells as well, reported by Zhou et al. (2000). It is not clear exactly what differential roles the contrast sensitive and non-sensitive BO cells play. They may work in a hierarchical or complemental manner as elements of BOWN computation. This intriguing issue should be elucidated by future research.

Interestingly, the activities of the hypothetical G-cells collectively create medial axis-like signals (Craft et al., 2007). The medial axis transform (MAT) in computer vision research is a shape representation that captures the local symmetry of the shape (Blum, 1973; Feldman & Singh, 2006). This suggests that border-ownership/figure-ground computation and shape computation processes are linked by the mechanism of BOWN computation proposed by Craft et al. through the dynamic interactions between BOWN sensitive neurons and G-cells (Craft et al., 2007; Feldman et al., 2013; Froyen et al., 2010). For example, Froyen et al. (Froyen et al., 2010) developed a Bayesian model which introduced a computation of medial axis (or “skeleton”) as a cue to border-ownership. In this model, BOWN signals developed dynamically as a result of competition between skeletons on both sides of the border. Furthermore, it has been argued that a BOWN computation mechanism underlies illusory contour perception (Kogo et al., 2010; Kumaran et al., 1996; Sajda & Finkel, 1995). The mechanism sketched above is in agreement with the hypotheses that IC cells are BO cells, and that BOWN computation is done by G-cells. This is also supported by neurophysiological evidence of neural coding of the medial axis in IT (Hung et al., 2012), and G-cell-like activities in response to illusory surface images in V4 (Cox et al., 2013). Although it is still speculative at this stage, it is possible that BOWN computation mechanisms mediate computational processes such as illusory contour perception, figure-ground organization, and shape perception, and play a pivotal role in the emergence of global properties in visual perception. Therefore, how exactly BOWN signals are computed in the visual system, and how they are linked to other neural processes for perceptual organization are vital pieces of information which need to be investigated further in future research.

In this paper, we presented a model that detects global coherency of BOWN signals based on the convexity relationship and consistency of surface properties. We showed that this approach gives extremely robust responses when reproducing human perception of complex images. The results strongly support the idea that large-scale consistency of surface properties is reflected in figure-ground organization through the interactions of the contrast-sensitive BO cells.

## Acknowledgement

Naoki Kogo was supported by a post-doctoral fellowship of Fonds voor Wetenschappelijk Onderzoek (FWO-Flanders post-doc grant 12L5115N, University of Leuven, 2014∼2017)

---

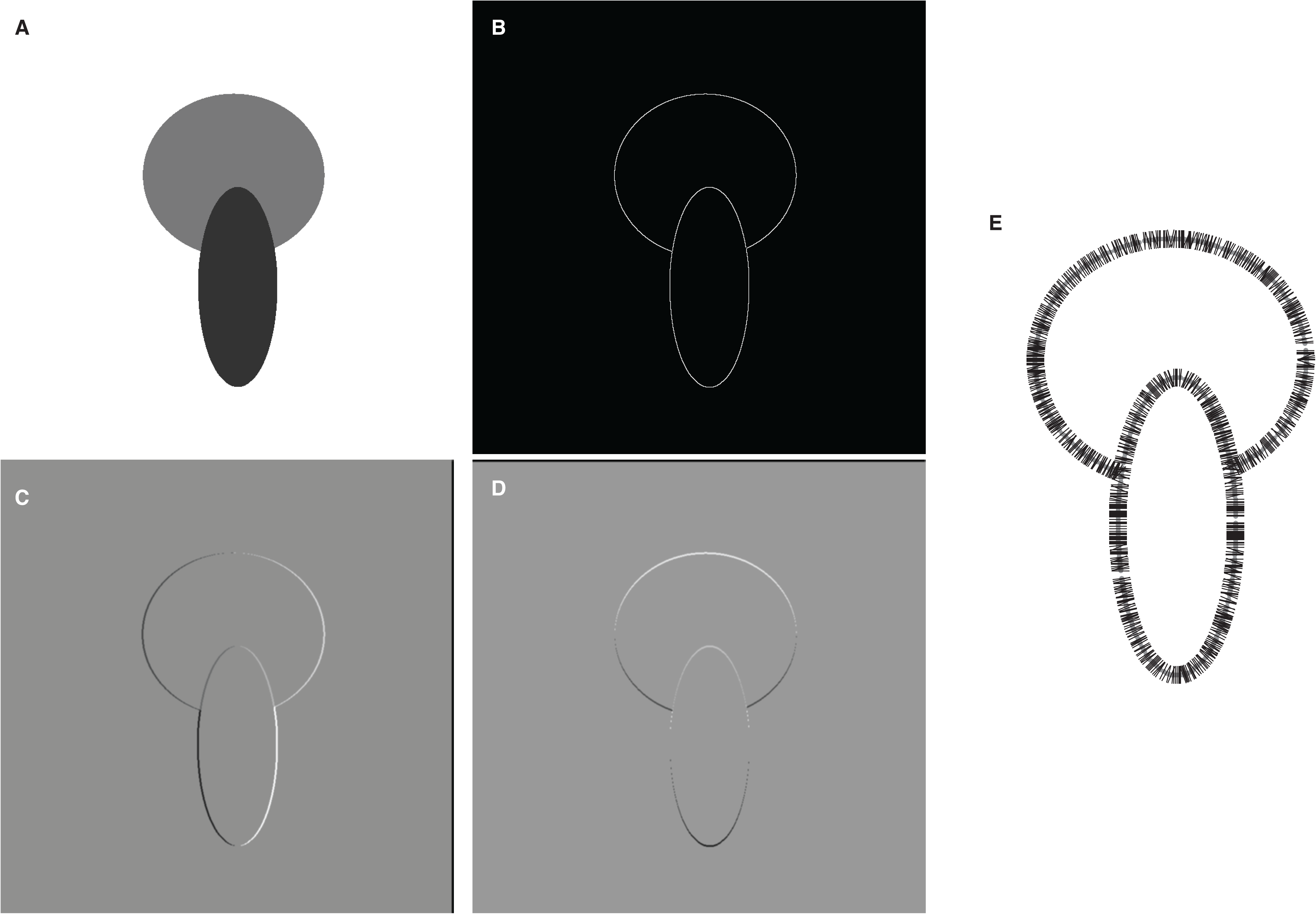

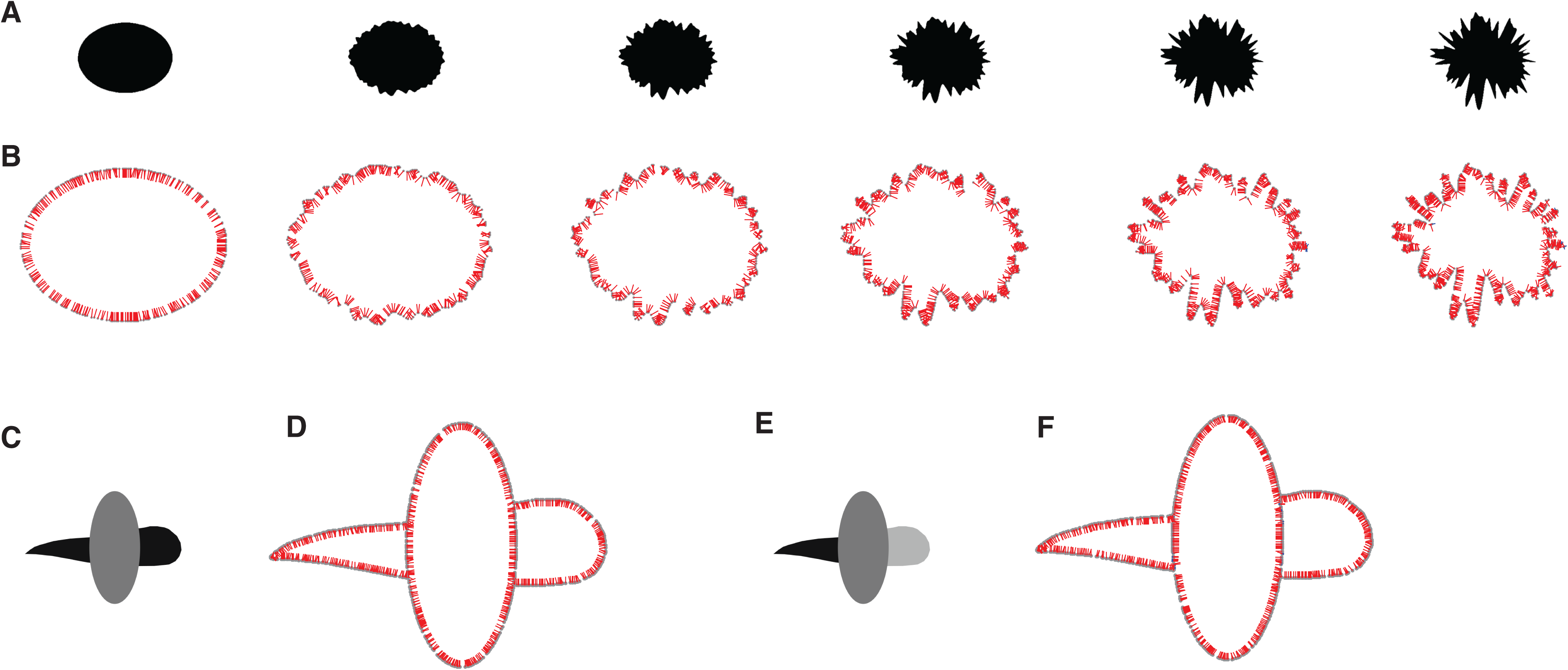

## Footnotes

1 We use the convention of using the terms convexity and concavity: A curvature of a contour of an area is called convex or concave by looking at it from outside of the area.

2 Note that the model does not allow for negative “activities”, hence if *b*^*t*^(*i*) < 0, then *b*^*t*^(*i*) = 0.

3 Note that number of iterations is different from the computation time. At each iteration, the interactions of all BOWN signals are computed. Hence, a higher number of iterations is not due to the need for more sequential processes of a higher number of BOWN signals. The number of iterations indicates how many repetitions of the processes are needed to settle on a final BOWN. Hence, this reflects the complexity and ambiguity of the images and, in turn, may correspond to the probability of establishing expected perceptions.

4 Although it has been reported that BO cells are also sensitive to stereo disparities (Qiu & von der Heydt, 2005), whether BO cells encode quantitative depth or not is unknown.

