## Supplement text for "Emergence of border-ownership by large-scale consistency and long-range interactions: Neuro-computational model to reflect global configurations"

### Supplemental material

#### S1. Pre-processing

The pre-processing stage of the model takes in an image and extracts from it edge orientations and surface properties, which are assigned to border nodes. Given a gray-scale image $A$ of size $I$ x $J$ (see for example Figure S1A) we first process the image through what we call color clustering. During this process, we compute the frequency of each gray-scale value. Those values that have a frequency below a certain threshold (in our case $0.005$ x $I$ x $J$ pixels) were discarded. Their corresponding pixels were then assigned the gray-scale value of the nearest pixel having a gray-scale value above the threshold. This procedure results in a new image $A^*$, which will lower the chances of getting spurious edges caused by slight luminance differences during the subsequent edge detection phase.

On this processed image $A^*$ we then run a Canny edge detector (Figure S1B) using the built-in MATLAB *edge* function, resulting in a set of edges $E = \{e_1 \ldots e_N\}$ (with each $e_n$ representing a two-dimensional vector in $R^2$, the location of the edge). In order to find their local orientations, we take the luminance gradient of $A^*$ in the X and Y directions, resulting in two gradient images $dX$ and $dY$ (Figures S1C and S1D). Given these, we have effectively defined the normals of the luminance gradient at each location in image $A^*$, and thus also the normal at each edge $e_n$:

$$N_x(n) = \frac{dX(i,j)}{|(dX(i,j), dY(i,j))|}$$

$$N_y(n) = \frac{dY(i,j)}{|(dX(i,j), dY(i,j))|}$$

where $i$ and $j$ define the location of edge $e_n$ in $A^*$. Given this information, we can now create two BOWN signals ($B_n$ and $B_{N+n}$) at each edge location, $e_n$, resulting in $2N$ BOWN signals (Figure S1E). Each competing pair then has the same location, but opposite normals (or BOWN directions):

$$B_n = \{e_n, N_x(n), N_y(n)\}$$

$$B_{N+n} = \{e_n, -N_x(n), -N_y(n)\}$$

Finally, for each BOWN signal $B_n$ gray-scale of the surface on the owned side (the side the BOWN signal indicates the ownership) and the un-owned side are detected. This was done by picking the gray-scale value in $A^*$ 4 pixels away along the BOWN signal direction, resulting in gray-scale vectors $C_{own}(n)$ and $C_{unown}(n)$.

**S2. Responses of the model to additional images**

In Figure S2 (A), a series of images are shown, created by systematically increasing the complexity of the contour of the oval (the leftmost image). This was done by first converting existing boundaries of a closed shape (oval) to a B-spline function. Subsequently, we added gaussian noise to the control points of this B-spline function in the direction of the normal along the boundary of the shape. Lastly, the new control points were used to generate a smooth noisy shape. The responses of the model (standard DISC2) are shown in (B). Because of the global interactions between the BOWN signals, the resulting BOWN is inward in all locations, despite the local complexity, indicating that the black area is considered to be figural, consistently along the border.

In Figure S2C and E, images modified from Yin, Kellman and Shipley (1997) and the responses of the model to them (D and F, respectively) are shown. In E, when the surface colors on the left and right areas are different, their colinear edges do not interact because of the inconsistent surface property. Nevertheless, the result shows that these areas are occluded by the central surface: the ownerships of the borders between the central area and the side areas belong to the central area.

#### FIGURE CAPTION

**Figure S1**: Illustration of pre-processing: (A) Input image; (B) Edge map retrieved by Canny Edge detection; (C) X and (D) Y direction gradients of luminance; (E) Sample set of resulting BOWN signals (black lines perpendicular to edges). Note that at each location, there is a pair of competing BOWN signals for two opposite-owner sides.

**Figure S2**: (A) Images modified from an oval (the leftmost image) by adding Gaussian noise on the contour to systematically increase the complexity of the shape. The mean of the random value was set to 100 pixels, and the standard deviation to 0, 5, 10, 15, and 20 pixels (from left to right). (B) Responses of the model showing inward ownership. (C-F) Images (C and E) modified from Yin, Kellman and Shipley (1997) and the responses of the model to them (D and F).
